# Transcriptional changes across tissue and time provide molecular insights into a therapeutic window of opportunity following traumatic stress exposure

**DOI:** 10.1101/2024.11.01.621484

**Authors:** Lauren A. McKibben, Meghna Iyer, Ying Zhao, Roxana Florea, Sophia Kuhl-Chimera, Ishani Deliwala, Yue Pan, Erica M. Branham, Sandrine M. Géranton, Samuel A. McLean, Sarah D. Linnstaedt

## Abstract

Unfortunately, survivors of traumatic stress exposure (TSE) frequently develop adverse posttraumatic neuropsychiatric sequelae (APNS) such as chronic pain and stress/depressive symptoms. Increasing evidence indicates that there is a ‘window of opportunity’ following TSE in which therapeutic interventions are most effective against APNS, yet mechanisms accounting for this observation are poorly understood. Here, we aimed to better understand such mechanisms by generating snapshots of the transcriptional landscape in the early aftermath of TSE across tissues and time. Adult rats were exposed to a TSE model, single prolonged stress (SPS). Then, eight tissues (hypothalamus, left and right hippocampus, amygdala, dorsal root ganglia, spinal cord, heart, and muscle) were isolated from these animals at 2, 24, and 72 hours after SPS and in unexposed controls (n=6 per group). mRNA expression from deep sequencing was used to identify differentially expressed genes (DEGs) and biological pathways enriched over time. In all tissues except the amygdala, the highest number of DEGs was observed 2-hours post-SPS, but DEGs were detected at all timepoints and in all tissues. Some transcripts were differentially expressed in a consistent manner across multiple tissues at a time point (e.g. *Fkbp5*, 2 hours post-SPS), while others had tissue-or region-specific expression patterns. Stress system pathways were most represented at 2-hours post-SPS, then stress/circadian/inflammatory pathways at 24-hours, and inflammatory pathways at 72-hours. Together these findings provide insights into post-TSE transcriptional landscape dynamics and suggest specific intervention windows of opportunity. Future validation is needed across sex, age, stressor, and cell type.

## Introduction

Almost all individuals will experience at least one traumatic stress exposure (TSE) in their lifetime^1^. While most people recover following TSE, a substantial proportion develop adverse posttraumatic neuropsychiatric sequelae (APNS) such as chronic pain, posttraumatic stress, and depressive symptoms^1–4^. APNS are often comorbid^4–7^ and incur tremendous societal costs^8^ including lost workdays^9^, decreased mental and physical health scores^10–13^, greater medical costs^9^, and increased incidence of suicidal ideation^14,15^. In addition, once APNS develop they are difficult to treat and can lead to opioid addiction or misuse^7,16–18^. Novel therapeutic strategies that prevent and/or treat APNS are urgently needed.

Results of previous research suggest that pharmacologic interventions which modulate the stress response may reduce APNS if administered in the early aftermath of TSE^19–31^. In animal models, early post-TSE intervention with propranolol^21,27,28^, minocycline^23^, high-dose corticosterone or hydrocortisone^22,24^, RU486^29^, morphine^19^, naloxone^21^, ketamine^20^, or SAFit2 (an FKBP51 inhibitor) ^25^ have been found to reduce APNS-like behavior. In several small human trials or observational studies, early post-TSE interventions have been found to improve or be associated with reduced APNS outcomes ^24^ ^26,30,31^.

Despite this evidence, relatively little is known regarding key molecular pathogenic mechanisms in the early post-TSE period. Such potential mechanisms include data suggesting that early post-TSE pharmacologic interventions may block emotional memory formation (e.g., estrogen^32^ or corticosterone/dexamethasone^33^) while other early interventions might hinder memory reconsolidation (e.g. propranolol^21,27^ and naloxone^21^). Increased understanding of the molecular changes occurring in the early aftermath of TSE will help provide insight into time-critical changes in the pathobiological mechanisms leading to APNS, thereby refining knowledge of mechanistic changes relevant to known therapeutic targets, defining the duration of the ‘window of opportunity’ and its specificity for certain targets, and promoting the identification of novel therapeutic targets for early post-TSE interventions.

One method of gaining insights into such changes is the evaluation of changes in RNA expression after TSE. RNA expression levels are representative of the underlying molecular biology of various phenotypes for several reasons: 1) RNA expression is responsive to environmental stimuli^34^, 2) their changes are easy to measure^34^, 3) RNA expression often reflects protein expression levels (which are fundamental drivers of biological processes)^35,36^, and their signatures can be extrapolated into biological pathways of importance^37,38^. Of relevance to the current study, multiple previous studies have identified specific RNA transcripts that change in response to TSE (e.g.^37–44^) or have performed large-scale re-analysis of combined datasets^45^. However, no previous studies have comprehensively examined RNA expression changes across multiple tissues and timepoints following TSE in a single study.

In the current study, we aimed to generate an overview of the transcriptional landscape following TSE by examining RNA expression changes across tissues and time, starting in the early aftermath of TSE and continuing through three days post-TSE. This timing enabled us to caputure acute transcriptional changes post-TSE. To simulate single-timepoint/large magnitude TSEs that occur in humans, we used a well-validated rodent model, the single prolonged stress (SPS) model. SPS exposes animals to multiple stressor types in a single day and has been shown to elicit a robust stress response^46^. Following SPS, we dissected eight tissues for analysis across the three post-SPS timepoints. We selected a subset of central nervous system and peripheral tissues for gene expression analysis because of their implication in APNS development (e.g. chronic pain and posttraumatic stress)^27,39,47–52^. In addition, a subset of key transcriptional findings were replicated across rodent species, labs, and RNA detection techniques.

## Materials and Methods

### Animals

All animal studies were completed in accordance with the Institutional Animal Care and Use Committee (IACUC) at the University of North Carolina at Chapel Hill (UNC-CH) under an approved protocol (#21.256). Adult male Sprague Dawley rats (postnatal day 65 (PD65)) were obtained from Charles River Laboratories (Wilmington, MA, USA) and pair-housed under a 12-hour light/dark cycle (lights on at 07:00) within the Division of Laboratory Animal Medicine at UNC-CH. All experiments were conducted under light conditions. Rats had *ad libitum* food and water access and were acclimated to the animal facility for seven days prior to handling. After facility acclimation and prior to stress exposure and/or tissue collection, animals were handled for twenty minutes each day for at least three days over the course of a week. A cohort of mice were used to replicate key findings of this study (*Supplementary Methods*).

### Single Prolonged Stress Protocol

The single prolonged stress (SPS) model is characterized by a series of stressors on a single day without direct tissue injury as follows: physical restraint for two hours (DC L120, Braintree Scientific, Inc., Braintree, MA, USA), forced swimming for 20 minutes (water temperature 22°C, depth 20 cm), recovery for 15 minutes, then exposure to diethyl ether (75mL in a glass dessicator) until general anesthesia is induced (generally < 7 minutes)^46,53^. SPS has also been shown by our group and others to cause enduring hyperalgesia^50,51,54–56^. Rats were randomly assigned to either the SPS-unexposed control group (n=6) or one of three SPS groups (2, 24, and 72 hours post-SPS groups; n=6 rats per group). SPS was conducted on six animals per day over three days with the start of SPS synchronized to 09:00 hours on PD80.

### Tissue collection

Dissections were conducted across four separate days, with the time of euthanasia for each group calculated in reference to the start of diethyl ether exposure (e.g. 2 hours post-SPS corresponded to the time between loss of consciousness from ether to the time of sacrifice). SPS-unexposed control rat samples were collected at the same time of day as all other groups (∼14:00 hours). Each rat was euthanized via live decapitation and eight tissues were collected within 20 minutes by the research team, including dorsal root ganglia (DRG; bilateral L4-6), spine (L4-6), heart (apex only), left gastrocnemius muscle, left and right hippocampus, left and right amygdala (pooled as one sample), and hypothalamus. We selected these tissues with key considerations in mind. Practically, following euthanasia RNA integrity begins to degrade, and we selected a subset of all tissues to reduce the negative impact of extending dissection time on tissue RNA quality. Acknowledging that we could only gather a subset of all relevant tissues, we prioritized those strongly implicated in APNS outcomes, as identified in the literature^27,39,47–52^.

Tissues were cut to ≤0.5 cm thickness and immediately placed in 5x volume RNAlater (Invitrogen Cat# AM7020, Waltham, MA, USA). After 5 minutes room temperature incubation, each sample was placed on ice (≤30 minutes) until it could be transferred to a -80°C freezer for long-term storage. Trunk blood was collected from all animals at the time of dissection, processed into serum, and stored on ice (≤30 minutes) before transfer to -80°C storage.

### Serum collection and processing

Whole blood collected from each animal was incubated at room temperature for 20 minutes to encourage blood coagulation and then phase separated by centrifugation at 3,000 x g for 10 minutes at room temperature. The top serum layer was transferred to a sterile 1.5 mL Eppendorf tube and stored at -80°C.

### Measurement of serum corticosterone levels

Serum corticosterone levels were measured using an enzyme linked immunosorbent assay (ELISA; Invitrogen Cat# EIACORT) according to the manufacturer’s instructions. Absorbance was measured at 450 nm using a Synergy HTX Multi-mode Microplate Reader (BioTek Instruments, Winooski, VT, USA).

### RNA isolation from animal tissues

5-10 mg of each tissue sample (24 animals x 8 tissues) were used for Trizol-based RNA extraction (Invitrogen Cat# 15596026) following the manufacturer’s standard protocol (*Supplementary Methods*). Samples were processed individually. After processing, RNA was preliminarily assessed for quality and quantity using a spectrophotometer (Nanodrop One, ThermoScientific, Waltham, MA, USA), then stored at -80 °C.

### mRNA Sequencing, alignment, and normalization

RNA sequencing was performed at Novogene Corporation Inc. (Sacramento CA, USA). All samples passed RNA integrity threshoholds and were library-prepped using Novogene’s custom protocol (additional details in the *Supplementary Methods*). More than 40 million raw reads were generated per sample. Samples were not pooled.

Raw sequencing reads were aligned to the Rat Genome Assembly mRatBN7.2 by using STAR 2.7.6a. Expression levels of each transcript were estimated via Salmon 1.4.0. Transcripts were filtered (*Supplementary Methods*) and the remaining read counts were normalized using DESeq2^57^

### Assessment of RNA expression patterns via principal components, differential gene expression analysis, and KEGG pathway analysis

To visualize clusters of closely related data points in our sequencing data, we performed principal component analyses^58^ on the full matrix of normalized read counts for each timepoint with RNA expression data (i.e. SPS-unexposed, 2, 24, 72 hours). The first two principal components were used to graph and visualize the largest variation in the data.

DESeq2 was used to assess gene expression changes over time following SPS relative to SPS-unexposed control gene expression. Gene transcripts were defined as differentially expressed genes (DEGs) if they met fold change and *p*-value thresholds set *a priori* (|log_2_[*fold change*]|≥1 and *p≤* 0.05). DEGs at each post-SPS timepoint within each tissue were visualized as volcano plots using GraphPad Prism (Boson, MA, USA). Common DEGs across tissues and timepoints were identified using an online list comparator tool (molbiotools.com).

### KEGG Pathway Analysis

We conducted Kyoto Encyclopedia of Genes and Genomes (KEGG) pathway analysis (ShinyGO v0.80^59^) using a prioritized list of DEGs within each tissue and at each timepoint whose [|log2*(fold change)*| × (-log *p*)] ≥ 1.

To better visualize changes over time in the top molecular pathways, and because many KEGG pathways shared similar gene constituents, we then conducted a blinded manual annotation of all KEGG pathways to assign one of ten broad mechnanistic categories to each pathway. Additional details of the KEGG analysis and manual annotation are in the *Supplementary Methods*.

Within the top KEGG category at each post-SPS timepoint, we then identified the top twelve DEGs based on the average of the gene expression product values [|log2*(fold change)*| × (-log *p*)] across all tissues at that timepoint. Finally, we visualized these top twelve DEG’s log2(*fold change*) across time and tissues as heatmaps using GraphPad Prism.

## Results

### Study design

As depicted in **Figure 1**, tissue was collected from SPS-unexposed control or SPS-exposed rats at one of three timepoints following SPS: 2 hours, 24 hours, and 72 hours (n=6 animals per group). From each animal at each timepoint, blood serum and eight tissues were collected (**Figure 1**). In a secondary cohort of age, sex, and strain matched rats, we collected serum from eight animals 3 minutes following SPS. In total 192 samples were collected from solid tissues and 32 samples from blood serum. RNA was extracted from each solid tissue sample and processed to generate RNA sequencing data. Corticosterone levels were measured from serum samples.

**Figure 1.**
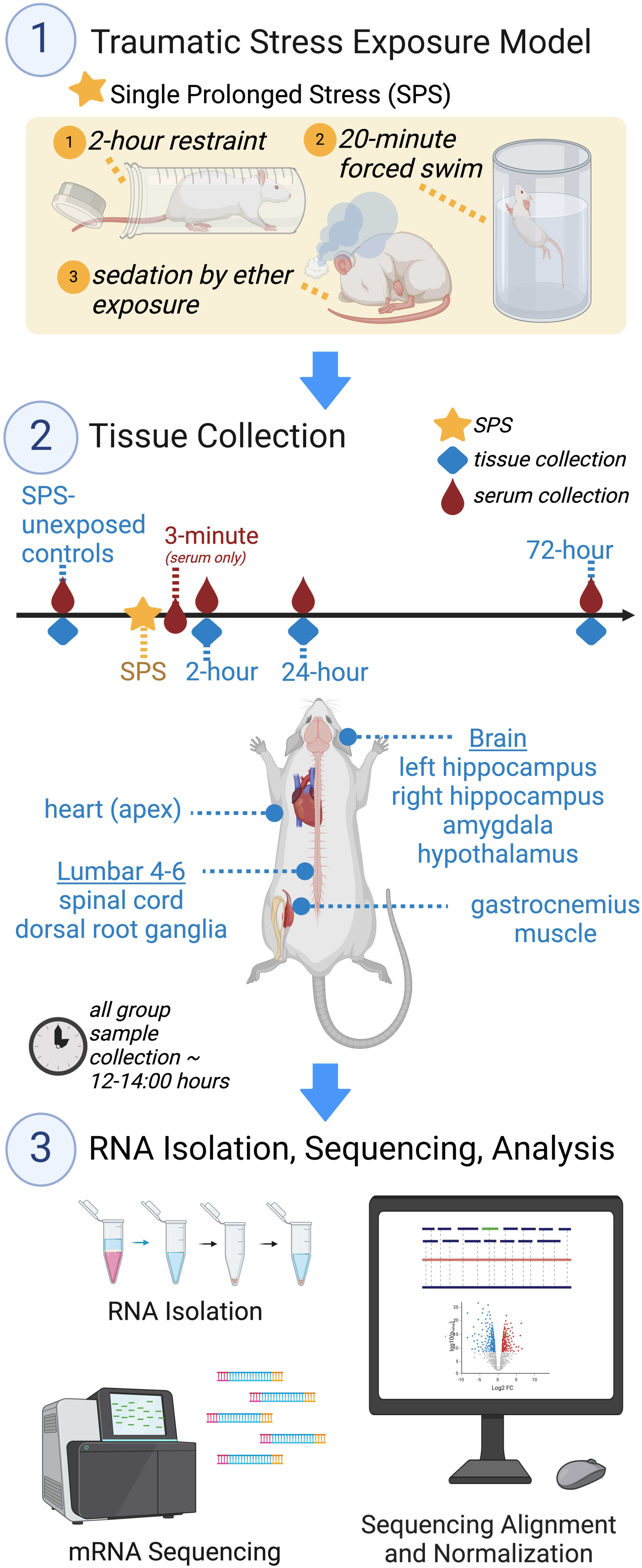
Experimental design and timeline. 1) The single prolonged stress (SPS) protocol was used to model traumatic stress exposure. This stress model is characterized by three multimodal stressors in a single day. 2) Tissues were collected for RNA expression studies from SPS-unexposed control rats and at three post-SPS timepoints: 2 hours, 24 hours, and 72 hours. At all timepoints, the following tissues were collected: left and right hippocampus, amygdala, hypothalamus, heart apex, lumbar (L) 4-6 spinal cord and dorsal root ganglia, and left gastrocnemius muscle. In addition, blood serum was collected at each of the 2-, 24-, and 72-hour timepoints as well as an additional 3-minute post-SPS timepoint. Blood serum samples were used for corticosterone measurements only (no RNA sequencing). 3) RNA was isolated from tissues and messenger RNA sequenced. Next, RNA expression data was aligned to the rat genome, cleaned, and assessed for differential expression relative to SPS-unexposed RNA expression levels. Created in BioRender. Linnstaedt, S. (2024) BioRender.com/s76h823

In adult (∼PD60) C57BL/6J mice, tissue samples were collected 2-hours following mSPS or from mSPS-unexposed mice. In another cohort of mice, blood plasma samples only were collected 3-minutes post-mSPS or in mSPS-unexposed control mice.

### Circulating corticosterone levels changed over time in response to single prolonged stress exposure

The purpose of the current study was to assess RNA expression changes over time in response to SPS across multiple tissues. However, prior to RNA based analyses, we first validated that SPS elicited a stress response in our animals via measurement of the stress hormone corticosterone. In rats, we found that serum corticosterone levels increased robustly 3 minutes following SPS (6.5x-fold increase relative to SPS-unexposed controls, t(20)=15.25, *p*<0.0001, **Figure 2A**). In mouse plasma collected 3 minutes post-mSPS, we also found an increase in corticosterone levels compared to mSPS-unexposed mouse serum collected 3 minutes post-mSPS (12.3x-fold increaste, t(8)=2.70, *p*=0.0271; *Supplementary Figure 1*). In addition, in rats we detected a slight decrease in serum corticosterone levels at the 2-hour timepoint following SPS relative to SPS-unexposed rats (2.1x-fold, t(24)=2.167, *p*=0.040, **Figure 2A**) but this decrease did not survive multiple hypothesis testing (Tukey’s adjusted *p*=0.310), nor did we detect such a decrease when we assessed plasma corticosterone levels in our 2-hour timepoint samples in mice following mSPS (1.1x-fold decrease, t(10)=0.641, *p*=0.5359, *Supplementary Figure 1*). Of note, corticosterone levels in the SPS-unexposed animals were consistent with previously published levels in adult male rats^54,60^ and mice^61^.

**Figure 2.**
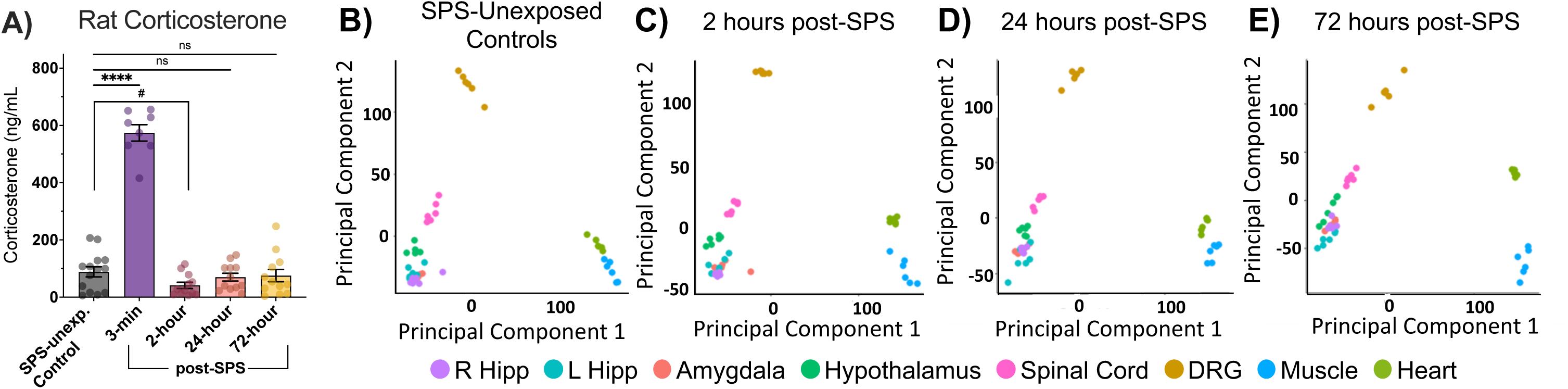
Blood levels of corticosterone measured in samples collected from rats at multiple timepoints following SPS and PCA plots of DESeq2 normalized RNA read counts. A) Serum corticosterone levels in samples collected from SPS-unexposed control rats and in samples collected from rats following SPS at four timepoints. Bars represent mean ± standard error of the mean. Then, DESeq2 normalized sequencing data from all tissues was analyzed using PCA in B) SPS-unexposed control rats and C) 2-hour, D) 24-hour, and E) 72-hour post-SPS-exposed rats. Each panel shows the eigenvalues for the top two principal components, on the x-and y-axes respectively, which cumulatively account for 51% of variability across the sequencing data. Each tissue is colored according to the legend. *\*\*\*\* p<0.0001, *p<0.05. Abbreviations: ns-nonsignificant, ng/mL-nanograms/milliliter, SPS-single prolonged stress, R hipp-right hippocampus, L hipp-left hippocampus, hypot-hypothalamus, DRG-dorsal root ganglion, PCA-principal components analysis*.

### Data quality assessments indicated high quality RNA, RNA sequencing results, and accuracy of tissue isolation

RNA was isolated from all tissues except serum. Based on both spectrophotometer and bioanalyzer readings, the RNA isolated from all 192 samples was of high quality and purity (*Supplementary Figure 2*A and 2B). The RNA sequencing method also yielded high quality data, with more than 96% reads mapping to the rat genome *(Supplementary Figure 2*C and 2D). Raw and normalized sequencing read files are publicly available on the UNC Dataverse via: https://doi.org/10.15139/S3/T4OW9K.

To ensure sequencing data quality, we performed principal component analyses of normalized RNA expression data from each timepoint (SPS-unexposed, 2, 24, and 72 hours post-SPS) to determine whether the samples clustered together based on tissue of origin. We found that the top principal component (“PC1”, **Figure 2B-E**) distinguished nervous system tissue samples (i.e. left and right hippocampus, amygdala, hypothalamus, spinal cord, and DRG) from non-nervous system tissue samples (i.e. heart and muscle), and the second principal component (“PC2”) distinguished DRG from central nervous system and peripheral tissues. Combined, PC1 and PC2 explained 51% of the variability in expression data. In addition to clustering of samples based on tissue of origin, we also observed slight changes in the relative position of these clusters over time (**Figure 2B-E**).

### Differential gene expression analyses identified distinct patterns of gene expression over time and across tissues

The percentage of significant DEGs (*p≤*0.05) that exhibited a high magnitude of differential gene expression (|log_2_(*fold change*)|≥1) at each timepoint and in each tissue is shown in **Figure 3A** (*blue and red bars*). The highest percentage of DEGs was detected 2 hours post-SPS in all tissues except for the amygdala. In the amygdala, the highest percentage of DEGs was detected 24 hours following SPS. Generally, across all timepoints, peripheral tissues (heart and muscle) exhibited the greatest percentage of DEGs. The spread of statistical significance (*p*-value) and log_2_(*fold change*) of individual DEGs for each tissue over time are shown as volcano plots in **Figure 3B**, with gene-labeled full resolution figures of each individual volcano plot provided in *Supplementary Figure 3*A-X. A complete list of DEGs including metrics on log_2_(*fold change*), *p*-value, and FDR-corrected *q*-value are provided in *Supplementary File 1*.

**Figure 3.**
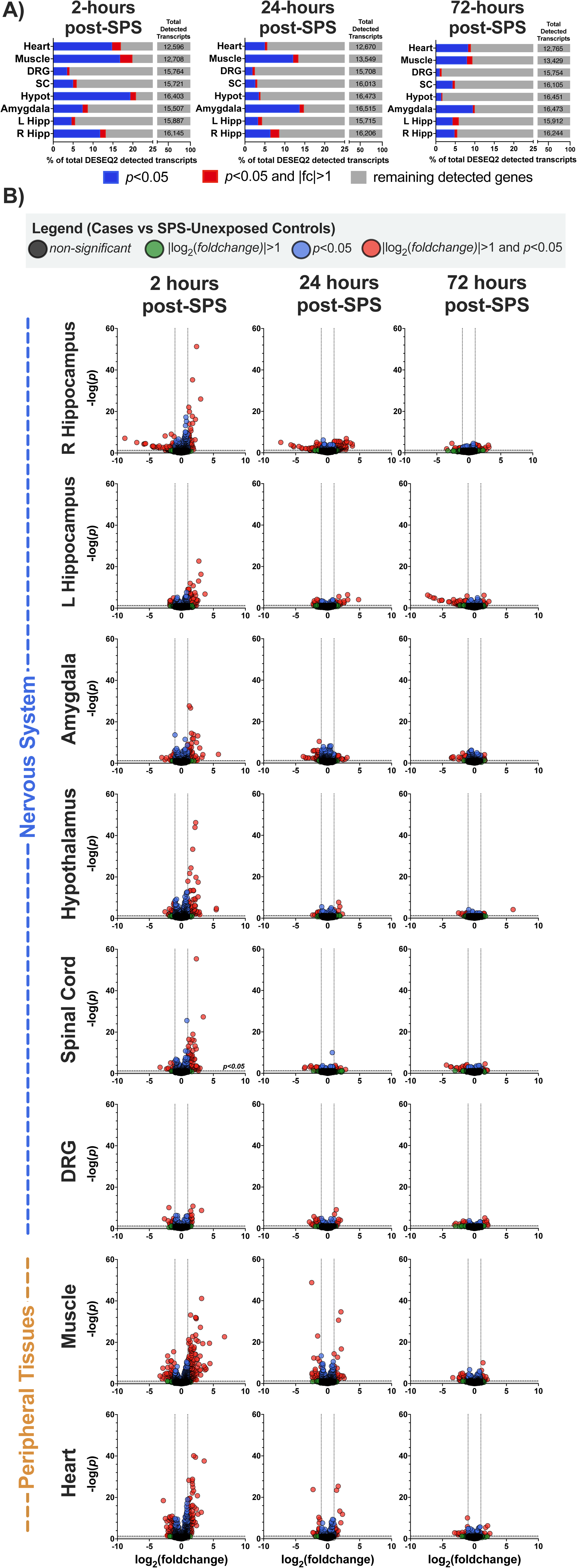
Overview of DEGs identified in each tissue and at each timepoint following SPS. A) The stacked bar graphs show percentages of DEGs out of all detected transcripts for each tissue and timepoint (total detected transcripts shown in right-most section of each bar). The red portion of each horizontal bar represents the percentage of DEGs that reach both thresholds: *p*≤0.05 and |log_2_(*fold change*)|≥1, the blue portion of each horizontal bar represents the remaining percentage of DEGs that only reach *p*≤0.05, and the grey portion represents the remaining gene transcripts that did not show differential expression. B) Each volcano plot shows the magnitude and statistical significance of DEGs within each tissue (rows) and at each post-SPS timepoint (columns). Tissues are separated into nervous system (top) and peripheral tissues (bottom). Negative log(*p*) is represented on the y-axis. A horizontal dashed line on each plot indicates the *p*=0.05 threshold. Log_2_(*fold change*) is shown on the x-axis. The vertical dashed grey lines on each plot indicate -1 and +1 log_2_(*fold change*) thresholds. Transcripts meeting *p*≤0.05 and |log_2_(*fold change*)|≥1 are represented by red dots, transcripts only passing *p*≤0.05 are represented by blue dots, transcripts only passing |log_2_(*fold change*)|≥1 are green dots, and transcript that met neither threshold are represented by black dots. The scale of the x and y axes are identical across all tissues and timepoints and were selected to include all dots. *Abbreviations: SPS-single prolonged stress, R Hipp-Right Hippocampus, L Hipp-Left Hippocampus, Hypot-Hypothalamus, SC-Spinal Cord, DRG-dorsal root ganglion, DEG-differentially expressed genes*.

Detailed volcano plots that include gene names (from *Supplementary Figure 3*) for nonbrain tissues are shown in **Figure 4**. Few of the top DEGs at the 2-hour post-SPS timepoint remained differentially expressed at the 24-and 72-hour post-SPS timepoints. We also show detailed volcano plots for brain tissues (**Figure 5**). Some transcripts were similarly altered by SPS across all brain tissues (e.g. *Gpd1* and *Mt1* at 2 hours post-SPS). However, the majority of DEGs were tissue specific (**Figure 5B**). Summary of a subset of these results including schematics indicating direction and magnitude of effect across specific genes in select tissue is shown in *Supplementary Figure 4*.

**Figure 4.**
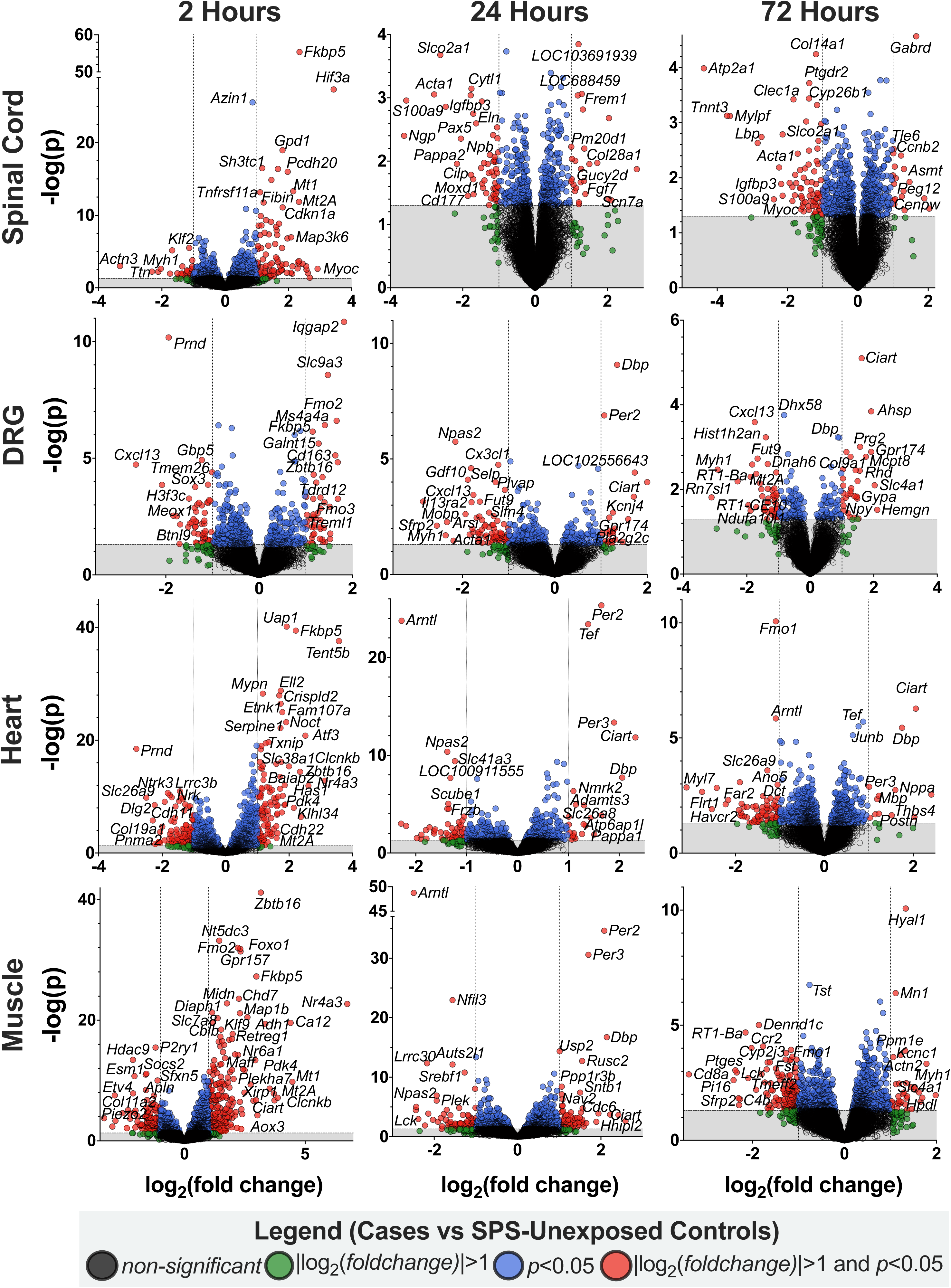
Non-brain tissue gene-transcript resolution of DEGs at all three timepoints following SPS. Volcano plots with gene-transcript names identified for several red dots. Note that axes are scaled to optimally show identities of DEGs rather than to match all axes (as in Figure 2). – Log(*p*) is represented on the y-axis. A horizontal dashed line on each plot indicates the *p*=0.05 threshold. Log_2_(*fold change*) is shown on the x-axis. The vertical dashed grey lines on each plot indicate -1 and +1 log_2_(*fold change*) thresholds. Individual DEGs are colored based on their *p*-value and *fold change* thresholds according to the legend. Of note, similar full resolution images of volcano plots for all tissues and timepoints is available in Supplementary materials. *Abbreviations: SPS-single prolonged stress, R Hipp-Right Hippocampus, DEG-differentially expressed genes*.

**Figure 5.**
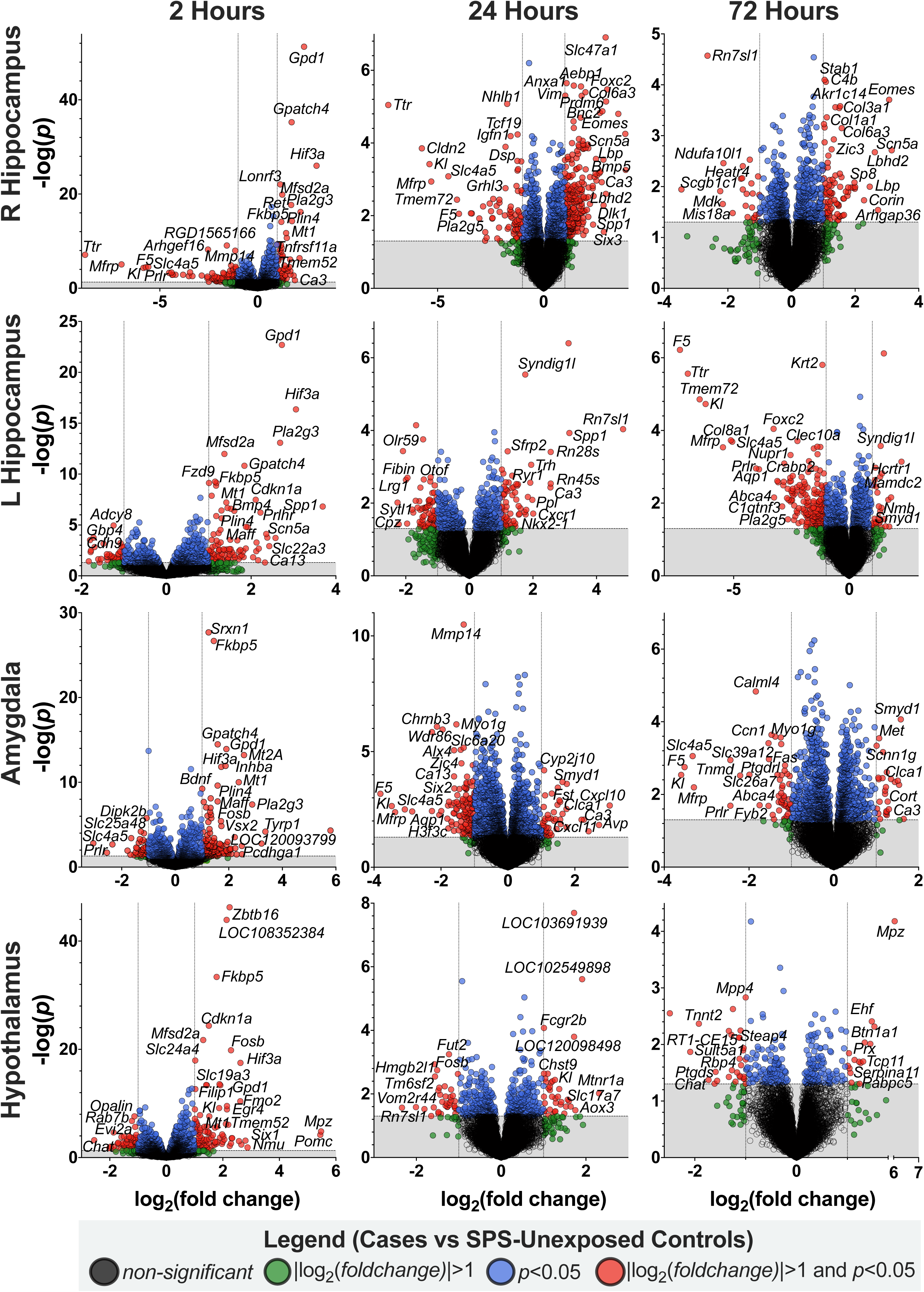
Brain tissue gene-transcript resolution of DEGs at all three timepoints following SPS. Volcano plots with top differentially expressed genes (DEGs) in the left and right hippocampus, amygdala, and hypothalamus at each post-SPS timepoint (2, 24, and 72 hours) as compared with SPS-unexposed animals. Note that axes are scaled to optimally show identities of DEGs rather than scaling to match all axes (as in Figure 2). –Log(*p*) is represented on the y-axis. A horizontal dashed line on each plot indicates the *p*=0.05 threshold. Log_2_(*fold change*) is shown on the x-axis. The vertical dashed grey lines on each plot indicate -1 and +1 log_2_(*fold change*) thresholds. Individual DEGs are colored based on their *p*-value and *fold change* thresholds according to the legend. Of note, similar full resolution images of volcano plots for all tissues and timepoints is available in Supplementary materials.

To further summarize gene expression changes following SPS, we next identified common and unique DEGs (*p*≤0.05 and |*log_2_(fold change)*|≥1) across all tissues and time points (*Supplementary File 2)*. Genes *Fkbp5* and *H3f3c* were robust and statistically significant DEGs in all eight tissues 2 hours following SPS (*Supplementary Figure 5*). By 24 hours, *Fkbp5* expression was no longer increased in any tissue while *H3f3c* remained decreased in expression across all tissues. Genes like *Zbtb16,* were significantly upregulated at 2 hours in most of the tested regions except the hippocampus or amygdala while other DEGs such as *Pla2g3* showed nervous system exclusive expression changes. No genes were commonly differentially expressed in all tissues at 72 hours. Overall, central nervous system tissues had more overlap in DEGs than did DRG, muscle, and/or heart (*Supplementary Figure 6*A-C). Within each tissue, most DEGs were unique to a specific time point, but some DEGs were significantly differentially expressed across all post-SPS timepoints (*Supplementary Figure 6*D-K).

### Validation of key changes in gene expression between SPS-unexposed and 2 hours post-SPS timepoints across labs using RT-qPCR (vs RNA sequencing), and in mice (vs rats)

The above RNA expression results were all based on RNA sequencing of tissues collected from rats. We validated some of the key DEG findings across labs, species, and RNA detection techniques by performing SPS in mice (mSPS), collecting tissues 2-hours following mSPS (and from mSPS-unexposed animals) and using RT-qPCR to quantify select transcripts. We selected transcripts *Fkbp5*, *Zbtb16*, and *Pla2g3* for validation. Besides *Pla2g3* in the heart, whose expression was not detectable in rats, we observed consistent expression patterns for all three transcripts 2-hours following mSPS in the hippocampus, hypothalamus, spinal cord, and heart (*Fkbp5* range of fold changes: 1.9-5.0x, *p* range: 0.0585-0.0003; *Zbtb16* range of fold changes:1.3-2.3x*, p* range*=*0.202-0.072; *Pla2g3* range of fold changes: 1.5-2.7x, *p* range*=0.380-0.0006; Supplementary Figure 7*), though the magnitude of this change in expression varied slightly.

### Pathway analyses identified specific KEGG pathways that were enriched in each tissue and timepoint following SPS

We next used DEGs identified from our RNA sequencing data to identify biological pathways enriched in each tissue and at each timepoint following SPS. All significant KEGG biological pathways at each timepoint and in each tissue are listed in *Supplementary File 3* along with their enrichment score, FDR *q-*value, product value, and assigned pathway categorization. To simplify data presentation, the overarching pathway categorization of the top 5 KEGG pathways (i.e. greatest [enrichment × -log(*q)*] value) for each timepoint and tissue are shown in **Figure 6A** (the full name, ID, and metrics of these top KEGG pathways are reported in *Supplementary Figure 8*). Among the top 5 pathways at each timepoint and tissue, Stress Signaling pathways were the most represented at 2 hours (n=15 KEGG pathways, Z=2.24, *p*=0.025 when compared to the category with the second highest KEGG pathway count). At 24-hours, Immune/Inflammatory pathways and Circadian pathways were the most common (n=7 KEGG pathways each, Z=2.63, *p*=0.008 when comparing both category counts combined with the next highest KEGG pathway count). Stress Signaling pathways (n=6 KEGG pathways) were still highly enriched, especially in the heart and muscle. At 72 hours, Immune/Inflammatory pathways were the most abundant (n=13 KEGG pathways, Z=2.50/2.76, *p*=0.012/0.006 when compared to the categories with the next highest pathway count).

**Figure 6.**
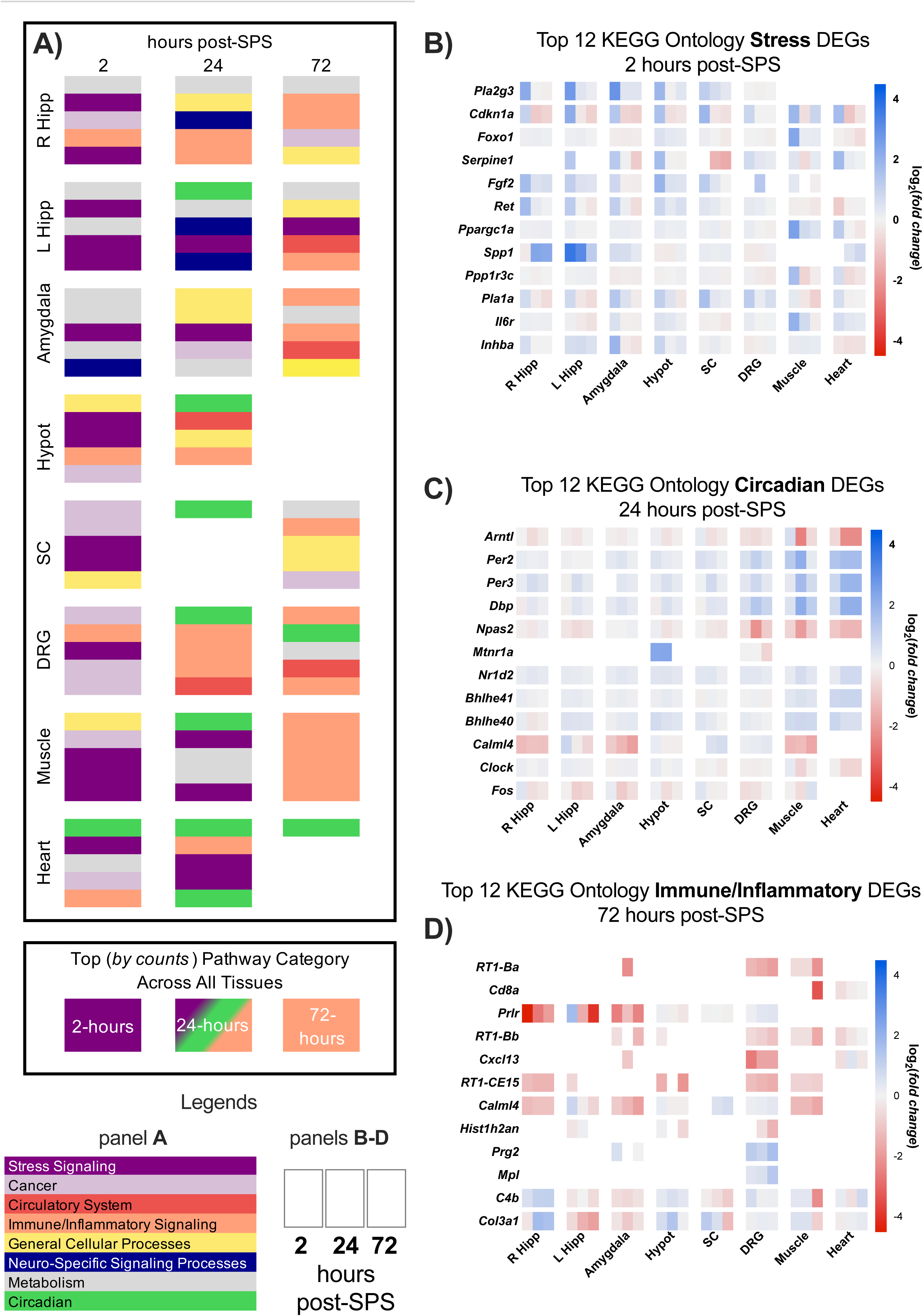
Summary of the most differentially expressed gene (DEG)-enriched biological pathways identified across all timepoints and tissues, and heatmaps of the top DEGs from the three most common pathway categories. A) (Top box) Kyoto Encyclopedia of Genes and Genomes (KEGG) pathways enriched in DEGs for each timepoint and tissue (FDR *q ≤* 0.1) were sorted in descending order based on |log_2_(*fold change*)| × -log(*p*). The categories of the top five KEGG pathways at each post-SPS timepoint and within each tissue are shown according to legend designations. Some tissues and timepoints had fewer than five enriched KEGG pathways, for example hypothalamus at 72 hours. (Bottom box) Summary of the most frequently identified biological pathways at each timepoint (as shown in top box). B-D) Heat maps of the log_2_(*fold change*) of the top twelve DEGs within each of the top three pathway categories identified in A: Stress Signaling, Circadian, and Immune/Inflammatory. Within each indicated tissue on the heatmaps, cells are ordered from left to right: 2 hours post-SPS, 24 hours post-SPS, and 72-hours post-SPS (as indicated in the legend). Positive fold-change values are in blue and negative fold-change values are in red. White spaces indicate that no expression was detected based on DESeq2 threshold parameters or no change in gene expression following stress was observed. *Abbreviations: R Hipp-Right Hippocampus, L Hipp-Left Hippocampus, Hypot-Hypothalamus, SC-Spinal Cord, DRG-dorsal root ganglion, FDR-false discovery rate*.

### Pathway-specific heatmaps provide pictorial details of longitudinal changes in gene expression following SPS

To illustrate the pattern of expression of the top gene transcripts belonging to each pathway identified in biological pathway enrichment analyses, we created heatmaps of the top 12 genes differentially expressed in the top KEGG pathway at each timepoint (**Figures 6B-D**). The log_2_(*fold change*) time course across all tissues for the top twelve 2-hour post-SPS gene transcripts within Stress Signaling KEGG pathways is shown in **Figure 6B**. Within this pathway, *Pla2g3* had the greatest average product score [|*log2(fold change)*| × (-log *p*)] across all tissues at 2 hours post-SPS (**Figure 6B**); this gene was exclusively expressed in nervous system tissues with the greatest *fold change* increase occurring at 2 hours post-SPS compared to SPS-unexposed control rats. We similarly identified the top twelve DEGs in Circadian pathways at 24 hours (**Figure 6C**) and Immune/Inflammatory Pathways at 72 hours (**Figure 6D**). These three heatmaps reveal both dynamic positive and negative differential gene expression over time and across different tissues within each of these overarching KEGG pathway categories. Although we found that each of these categories was highly represented at specific post-SPS time points, for many genes there was robust differential gene expression across several time points.

## Discussion

We used a well-validated rodent model to map the time course of gene expression changes within key brain and body regions after a substantial stressor. These central (hippocampus, amygdala, hypothalamus, spinal cord, DRG) and peripheral (heart, muscle) tissues/regions were chosen because together they play a dominant role in mediating acute and persistent phenotypic changes^49,62–64^. Gene expression changes in peripheral tissues 2 hours post-TSE were the most robust, but expression differences persisted in all tissues through 72 hours. Pathway-level analyses indicated that broad changes in the transcriptional landscape following TSE progressed from activation of stress system pathways in the early aftermath of TSE (2 hours), to a mixture of stress, circadian, and immune signaling pathways at 24 hours, to a predominance of immune/inflammatory pathways at 72 hours. In addition to these pathway-level data, we also identified gene transcripts that were distinctly activated at specific timepoints and tissues. For example, two genes, *Fkbp5* and *H3f3c,* were significantly altered in all tissues 2 hours post-SPS, while other genes such as *Mtnr1a*, *Grin2a*, and *Cxcl13* were up or down regulated in only one tissue. Additional observations from our analyses included substantial differences in gene expression following TSE in the left vs right hippocampus, a high degree of DEG overlap in central nervous system tissues, and <10 gene transcripts in each tissue that were consistently differentially expressed across all timepoints. Overall, this large-scale RNA sequencing data provides insights into the changing transcriptional landscape following TSE, provides information regarding mechanisms of the pathogenesis of APNS, helps in the identification of promising therapeutic targets for the reduction of APNS, and provides insight into a potential ‘windows of opportunity’ for secondary preventative interventions.

The agreement between past literature and our key findings bolsters confidence in the presented data and also provides replication of previous studies. For example, previous studies have shown a tightly controlled relationship between key mediators of the HPA-axis, such as between corticosterone levels and expression of the glucocorticoid receptor gene *Nr3c1* and a negative regulator of the HPA-axis *Fkbp5*^65–67^. We also observed a consistent relationship between these stress mediators: corticosterone levels were positively correlated with *Nr3c1* expression, and both corticosterone and *Nr3c1* levels were inversely correlated with *Fkbp5* expression. Further, a recent study showed that glucocorticoids induce expression of the neurogenic transcription factor *Zbtb16*^68^, and similarly we observed that *Zbtb16* is strongly induced 2 hours following SPS (in five of eight tissues analyzed). We also observed subsequent increased expression of *Eomes,* a gene that has been shown to partially mediate the neurogenic effect of *Zbtb16* ^68^, 24 and 72 hours following SPS in the right hippocampus.

Observed changes in gene expression at both individual transcript and pathway levels over time following TSE provide insight into potential mechanisms important to APNS pathogenesis. Stress, circadian, and immune systems are known to be substantially affected by TSE and are associated with APNS development (reviewed in ^52,69–71^). This study provides important new information regarding the timing, location, and evolution of these pathway activations. A more comprehensive, detailed understanding of these inter-relationships is essential to formulating and testing specific hypotheses (e.g., identification of key genes/proteins whose inhibition or augmentation prevents phenotypic changes, and the duration of pathway-specific secondary preventive therapeutic windows).

Part of the impetus for the current study was the observation by our team and others that promising therapeutic candidates for the reduction in APNS appear to be most effective at reducing APNS-like behaviors if administered early following TSE^19–31^. The current data provides specific molecular insights regarding why this might be the case. For example, we previously showed that a small molecule inhibitor of FKBP51 (the protein encoded by *Fkbp5*), SAFit2, only produces enduring reductions in pain-like behaviors if administered within 2 hours of TSE^25^. RNA expression data from this study indicate that *Fkbp5* mRNA expression levels increase dramatically 2 hours following TSE but return to baseline levels in all tissues by 72 hours. Given that protein translation dynamics generally parallel gene expression ^35,36^, these study data explain our finding regarding the optimal therapeutic timing of administering this protein inhibitor. The study data highlight the tremendous promise of this work for identifying different therapeutic targets, their timing of appearance/evanescence after TSE, and effective therapeutic interventions for novel targets.

Strengths of this study include a comprehensive assessment of gene expression changes across multiple timepoints and tissues, execution of high-quality RNA sequencing, control experiments confirming high quality RNA expression data and validation of tissue origin, and validation of key findings across labs, animal species, and method of detection. Nevertheless, this study also had several limitations. First, we focused this initial study of RNA expression following TSE on a single sex, single age, and single stressor. It is important to note that gene expression following TSE has been shown to differ according to factors such as sex, age, and stress type/magnitude (e.g. multiple stressors). Thus our results might not be generalizable to other animals and conditions. Second, we performed bulk RNA sequencing of RNA isolated from all cells in each indicated tissue. While such population-level work is informative and can lead to increased insight into post-TSE molecular mechanisms, previous studies have shown distinct gene expression differences across cell types within each tissue, suggesting that future studies would benefit from increased resolution of gene expression data according to cell type. Third, we did not consider individual variability in susceptibility and resilience to SPS and how these differences might influence gene expression, despite a previous study showing that individual genetics in outbred populations are likely to contribute to variability in mechanisms and behavior^72^. Fourth, we did not assess gene expression changes prior to 2 hours following TSE or beyond 72 hours following TSE. Therefore, it is uncertain whether any of the observed gene expression changes develop sooner or persist longer. Finally, pathway analyses were limited by the fact that many genes identified as DEGs in our study were not listed in the KEGG database, resulting in pathway results based on limited datasets (e.g. *Fkbp5*, which is known to play a role in a variety of pathways such as NFkκB, Akt, and steroid pathways, was only listed in the estrogen signaling pathway in KEGG).

In conclusion, we generated a comprehensive reference dataset of gene expression changes over time following TSE. Such information could help provide insights into the dynamic molecular signatures driving APNS and could lead to novel therapeutic insights into timing-related changes following TSE.

## Acknowledgements

Research reported in this publication was supported by the National Institute of Neurological Disorders and Stroke of the National Institutes of Health under Award Number R01NS118563 (Linnstaedt and McLean). Additional funding was provided by the Rita Allen Foundation (Linnstaedt), and the National Institute of General Medical Sciences under Award Number T32GM008450X (T32 Fellow: McKibben). The authors would like to acknowledge Brittania Wanstrath and Liz Albertorio-Saez for their help with animal tissue collection and Taanvii Verma for her help with figure revisions.

## Data Availability Statement

We provide raw and analyzed RNA sequencing data for public use, freely available at the following link: https://doi.org/10.15139/S3/T4OW9K. Other data generated as a part of the analyses presented in this mansuscript are available in Supplementary Information.

## Author Contributions (CREdiT Statements)

**Lauren A. McKibben:** Conceptualization, Methodology, Formal Analysis, Investigation, Data Curation, Writing-Original Draft, Writing - Review & Editing, Visualization, Supervision, Project Administration **Meghna Iyer**: Writing Original Draft, Writing - Review & Editing, Visualization **Ying Zhao**: Programming, Formal Analysis, Data Curation, **Roxana Florea:** Validation, Formal Analysis, Writing-Original Draft, Writing - Review & Editing, Visualization **Sophia Kuhl-Chimera**: Investigation, Writing - Review & Editing, Visualization **Ishani Deliwala**: Investigation, Writing - Review & Editing **Yue Pan**: Programming, Formal Analysis **Erica M. Branham**: Conceptualization, Methodology, Writing-Review & Editing **Sandrine M. Géranton**: Methodology, Validation, Writing-Original Draft, Writing - Review & Editing **Samuel A. McLean**: Conceptualization, Resources, Writing - Review & Editing, Funding Acquisition **Sarah D. Linnstaedt**: Concepturalization, Methodology, Resources, Writing-Original Draft, Writing - Review & Editing, Supervision, Project Administration, Funding Acquisition

