## Supplementary Tables and Figures for "Transcriptional changes across tissue and time provide molecular insights into a therapeutic window of opportunity following traumatic stress exposure"

#### Materials and Methods

##### *RNA isolation from animal tissues*

Brain tissue (including the left and right hippocampus, amygdala, and hypothalamus), spine, and DRG tissue were homogenized in Trizol using a bead mill homogenizer (5 meters/second, 20s). Heart and muscle tissues were homogenized by pestling (Cat# Axygen PES-15-B-SI, Corning Inc., Corning, New York, USA).

##### *mRNA Sequencing, alignment, and normalization*

The following experimental steps were performed at Novogene Corporation Inc. (Sacramento CA, USA). RNA quantity and quality of all samples was assessed using a BioAnalyzer 2100 (Agilent, Santa Clara, CA, USA). Only samples with more than 200 ng RNA and an RNA integrity (RIN) value above 6.0 could be sequenced. All 192 samples passed these thresholds. Libraries were prepared using Novogene's custom protocol as follows: messenger RNA (mRNA) was purified from total RNA using poly-T oligo-attached magnetic beads, mRNAs were fragmented and primed for reverse transcription via random hexamers, underwent end repair and A-tailing, were ligated to adapters, size selected, then amplified via polymerase chain reaction. The completed libraries were assessed for size, quality, and quantity using a Bioanalyzer and Qubit. Libraries were sequenced and multiplexed to obtain 150 nucleotide paired-end reads using an Illumina's NovaSeq 6000 system. More than 40 million raw reads were generated per sample. All samples passed a sequencing base-quality Q-score threshold of Q30, indicating high base-calling accuracy.

##### *Sequencing alignment and normalization*

Raw sequencing reads were aligned to the Rat Genome Assembly mRatBN7.2 by using STAR 2.7.6a. Expression levels of each transcript were estimated via Salmon 1.4.0. Raw read counts across all samples, regardless of tissue or timepoint, were filtered to remove genes with an average read count <1. Then, within each tissue, transcripts with an average read count <5 were also filtered out independent of timepoint. The total number of detected transcripts ranged from 12,596 to 16,515 across tissues. These remaining raw read counts for all samples were normalized using DESeq2[52] (i.e. normalization based on the geometric mean of gene counts across all samples).

##### *Assessment of RNA expression patterns by differential gene expression analysis*

DESeq2 was used to assess gene expression changes over time following SPS relative to SPS-unexposed control gene expression. For this analysis, read counts from each tissue at the 2-hour timepoint following SPS was compared to read counts from corresponding tissues from the SPS-unexposed control animals. We then repeated this analysis to compare read counts from the 24- and 72- hour timepoints, each to the SPS-unexposed control animals.

##### *KEGG Pathway Analysis*

In order to identify molecular pathways that were enriched in DEGs based on the most significant *and* strongest magnitude fold change, we calculated the product of -log(*p*-value) and |$\log_2$*(fold* *change)*| and conducted Kyoto Encyclopedia of Genes and Genomes (KEGG) pathway analysis (ShinyGO v0.80[54]) using this prioritized list of DEGs within each tissue and at each timepoint whose $[|\log_2(\textit{fold change})| \times (-\log p)] \geq 1$.

KEGG pathways passing a threshold of FDR $q \leq 0.1$ were identified for each post-SPS timepoint and in each tissue. Similarly to the product calculation used to identify genes with both high significance *and* robust fold change magnitudes, we used the product of each KEGG pathway enrichment score and its -log(*q*-value) to identify the top KEGG pathways. To better visualize changes over time in the top molecular pathways, and because many KEGG pathways shared similar gene constituents, we also blind-sorted the entire KEGG pathways database (KEGG.jp). Using our knowledge of the field and based on KEGG's existing categorization of their pathways, we sorted each pathway into one of ten overarching molecular process categories: Metabolism, Cancer, Circulatory System, Immune/Inflammatory Signaling, General Cellular Processes, Neuro-Specific Signaling Processes, Circadian Signaling, Genetic/Epigenetic Machinery, and Other. For pathways where categorization was not clear, we based our sorting on a brief literature search of the pathway and on its similarity in gene constituents with our ten molecular process categories. Within the top 5 KEGG pathways at each timepoint and within each tissue, we counted the occurrence of each category and conducted two-sample proportion tests to identify the top category(s) across all tissues at each timepoint. This enabled us to identify broad, time-specific changes in the transcriptional landscape.

Within the top KEGG category at each post-SPS timepoint, we then identified the top twelve DEGs based on the average of the gene expression product values $[|\log_2(\textit{fold change})| \times (-\log p)]$ across all tissues at that timepoint. Finally, we visualized these top twelve DEG's $\log_2$(*fold change*) across time and tissues as heatmaps using GraphPad Prism.

***Replication of key molecular findings in a mouse SPS model of TSE***

Animals. All mouse experiments were carried out in accordance with the guidelines of the UK Animals (Scientific Procedures) Act 1989 at University College London, UK (license #PP0091160). Male C57BL/6J mice were purchased form Charles River UK, at PD56 and were allowed a minimum of 7 days acclimatization before the start of experiments. Animals were housed in groups of 3-4 in a temperature- and humidity-controlled environment (20±2°C, 45-65%), with lights on from 07:00-19:00, and were given water and food *ad libitum*. All animals within a cage were assigned to either the control or experimental group. *SPS model (mSPS).*The stress paradigm was conducted between 09:00-13:00 hours and it consisted of sequential exposure to the following stressors as described previously[55] (adapted to use isoflurane instead of ether to adhere to UK animal research guidelines): physical restraint for one hour (Plexiglas tubes: 11x3cm LxD), forced swimming for 3 minutes (water temperature 23°C, depth 25cm), exposure to rat urine odour for 15 minutes (Envigo, Code S.T. - 0109), and lastly, 7 minutes exposure to 1% isoflurane mixed in $O_2$, flow rate 1.5L/min. Control mice were left undisturbed in their home cages.

*Tissue collection and RT-qPCR.* 2 hours following mSPS, mice were terminally anaesthetized with rising concentration of $CO_2$. Tissues (left and right hippocampus, hypothalamus, spinal cord lumbar [L4-6], and heart [apex]) were quickly collected and stored in RNAlater at -80°C until further processing. Total RNA was extracted using an acid phenol extraction method (QIAzol lysis reagent, #79306; RNeasy mini-kit, #74104; Qiagen, Hilden, Germany) previously described[56]. Samples were incubated with DNase I (#79254, Qiagen) and RNA concentrations were determined from Nanodrop measurements (Labtech International, Ringmer, UK). Equal amounts of RNA were reversed transcribed with a mix containing oligo $dT_{15}$

primer (C110A, Promega, Madison, WI, USA), Random Nanomers (R7647, Sigma-Aldrich, St. Louise, MO, USA), dNTP mix (U151B, Promega), and SuperScript III Reverse Transcriptase (#18080-093, Invitrogen) for 5 min at 25°C, 50 min at 50°C, and 15min at 70°C. The 1/10 cDNA samples were later prepared for RT-qPCR using SYBR Green JumpStart (S4438, Sigma-Aldrich) and the respective primers (*Supplementary Table 1*). Reactions were performed in triplicate and control cDNA samples without reverse transcriptase or cDNA template were included each time.

Corticosterone assay. Blood samples were collected in paediatric tubes (#459075, Greiner, Kremsmünster, Austria) between 10:00-15:00 hours (just before tissue collection) and kept on ice for 2 hours. Samples were then centrifuged for 10 minutes at 10,000 rpm, 4°C, and plasma was collected and processed using a Corticosterone ELISA kit (Ab108821, Abcam, Cambridge, UK) following the manufacturer protocol.

***Supplementary Tables and Figures***

| Oligo | Sequence (5' – 3') |
|---|---|
| *Fkbp5* Forward | CAATGCTGAGCTTATGTACG |
| *Fkbp5* Reverse | CTTTTCTTGGTGTCCATCTC |
| *Hprt* Forward | AGGGATTTGAATCACGTTTG |
| *Hprt* Reverse | TTTACTGGCAACATCAACAG |
| *Pla2g3* Forward | GCTCCCATACCCGTCACTG |
| *Pla2g3* Reverse | CCTGTGGTGTGTATCCCAGA |
| *Zbtb16* Forward | AGGACCGTAAGGCTCGATACC |
| *Zbtb16* Reverse | ACACTGGCGTAGCCACTCT |

***Supplementary Table 1.*** Primer sequences used in RT-qPCR to assess gene expression across mouse tissues collected 2 hours following mSPS exposure. *Abbreviations: mSPS- mouse single prolonged stress.*

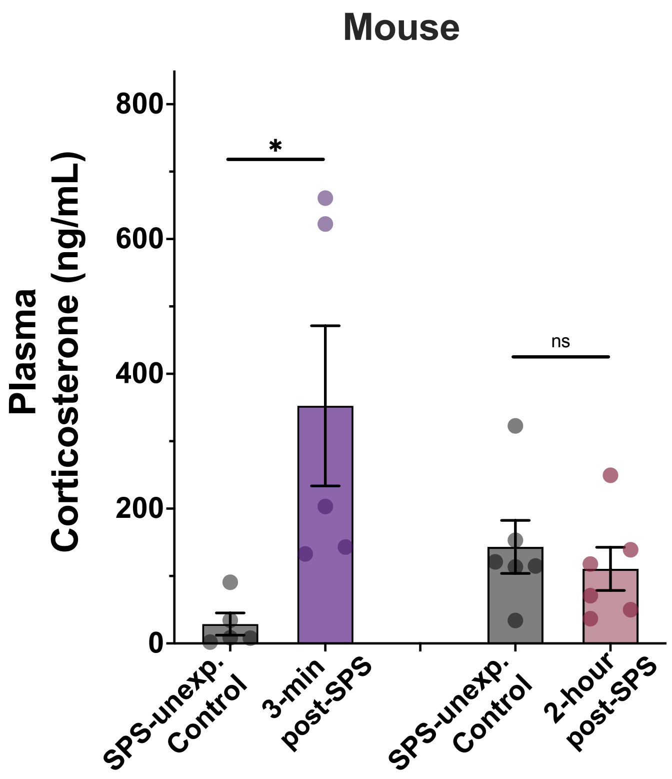


***Supplementary Figure 1.*** Blood levels of corticosterone measured in samples collected from mice at multiple timepoints following single prolonged stress (SPS). Serum corticosterone levels are shown in samples collected from mice that were either unexposed to SPS or exposed to SPS and samples collected 3-minutes or 2-hours later. Bars represent Mean ± SEM.*$^*p<0.05$. Abbreviations: ns-nonsignificant, ng/mL-nanograms/milliliter.*

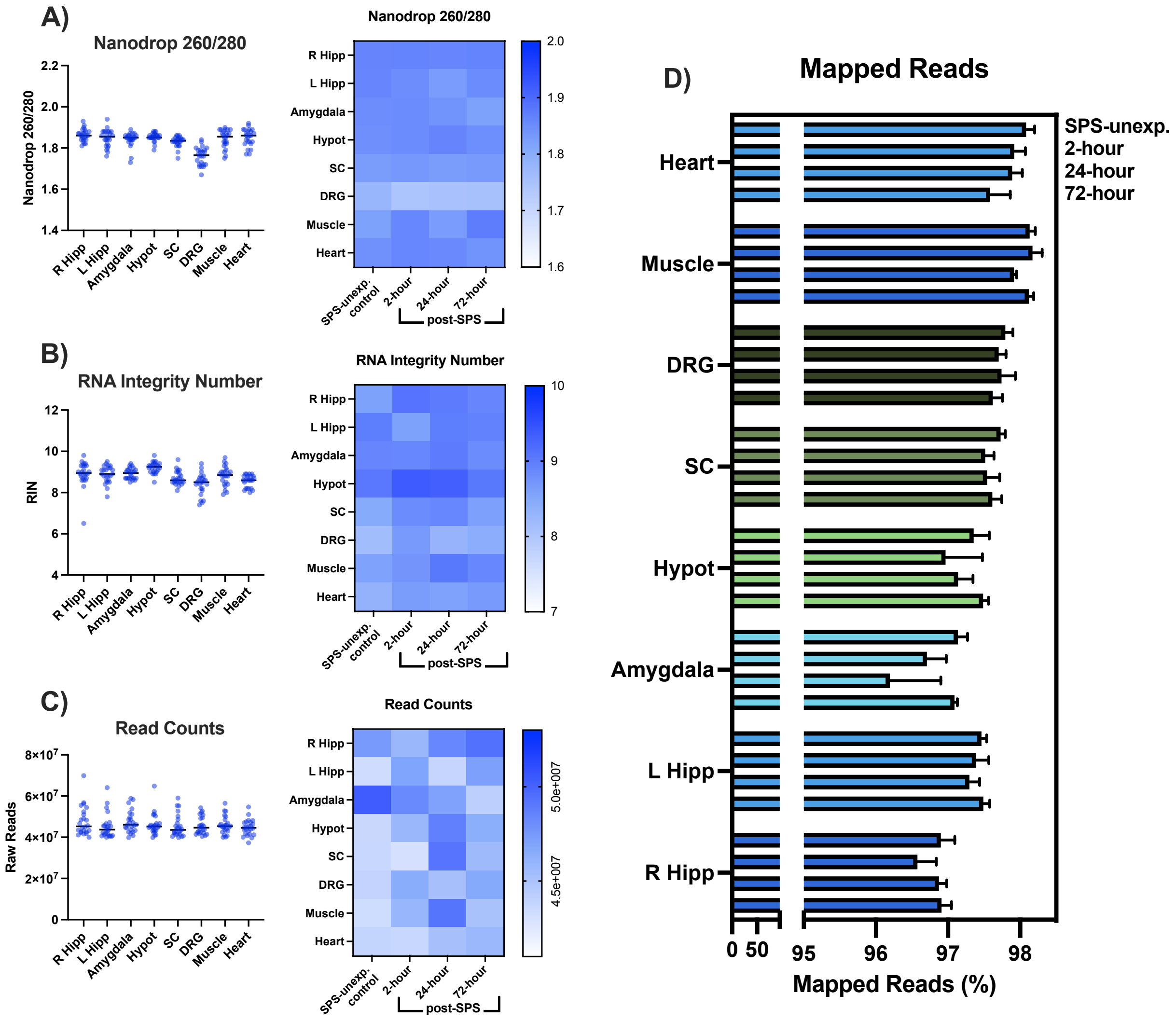


***Supplementary Figure 2****.* RNA and Sequencing Quality Metrics. Prior to mRNA sequencing, tissue-isolated total RNA was tested for quality using two metrics: A) 260/280 ratio, B) RNA integrity number. Following DESeq2 normalization, sequencing quality was assessed via: C) Total number of read counts and D) percentage of mapped reads across time and tissue. *Abbreviations: R Hipp-right hippocampus, L Hipp-left hippocampus, Hypot-hypothalamus, SC-spinal cord, DRG-dorsal root ganglion,, SPS-single prolonged stress, unexp.-unexposed.*

***Supplementary Figure 3.*** Each volcano plot from Figure 3B is shown (A-X) in large scale and full resolution with select gene ID labels. On each plot, negative Log(*p*) is represented on the y-axis. A horizontal dashed line on each plot indicates the $p=0.05$ threshold. $Log_2$(*fold change*) is shown on the x-axis. The vertical dashed grey lines on each plot indicate -1 and +1 $log_2$(*fold change*) thresholds. Transcripts meeting $p \leq 0.05$ and $|log_2(\textit{fold change})| \geq 1$ are represented by red dots, transcripts only passing $p \leq 0.05$ are represented by blue dots, transcripts only passing $|log_2(\textit{fold change})| \geq 1$ are green dots, and transcript that met neither threshold are represented by black dots.

*S3. A. Right Hippocampus 2 Hours*
*S3. B Right Hippocampus 24 Hours*
*S3. C Right Hippocampus 72 Hours*
*S3. D Left Hippocampus 2 Hours*
*S3. E. Left Hippocampus 24 Hours*
*S3. F. Left Hippocampus 72 Hours*
*S3. G. Amygdala 2 Hours*
*S3. H. Amygdala 24 Hours*
*S3. I. Amygdala 72 Hours*
*S3. J. Hypothalamus 2 Hours*
*S3. K. Hypothalamus 24 Hours*
*S3. L. Hypothalamus 72 Hours*
*S3. M. Spinal Cord 2 Hours*
*S3. N. Spinal Cord 24 Hours*
*S3. O. Spinal Cord 72 Hours*
*S3. P. Dorsal Root Ganglia 2 Hours*
*S3. Q. Dorsal Root Ganglia 24 Hours*
*S3. R. Dorsal Root Ganglia 72 Hours*
*S3. S. Muscle 2 Hours*
*S3. T. Muscle 24 Hours*
*S3. U. Muscle 72 Hours*
*S3. V. Heart 2 Hours*
*S3. W. Heart 24 Hours*
*S3. X. Heart 72 Hours*

**S3. A. Right Hippocampus 2 Hours Post-SPS**

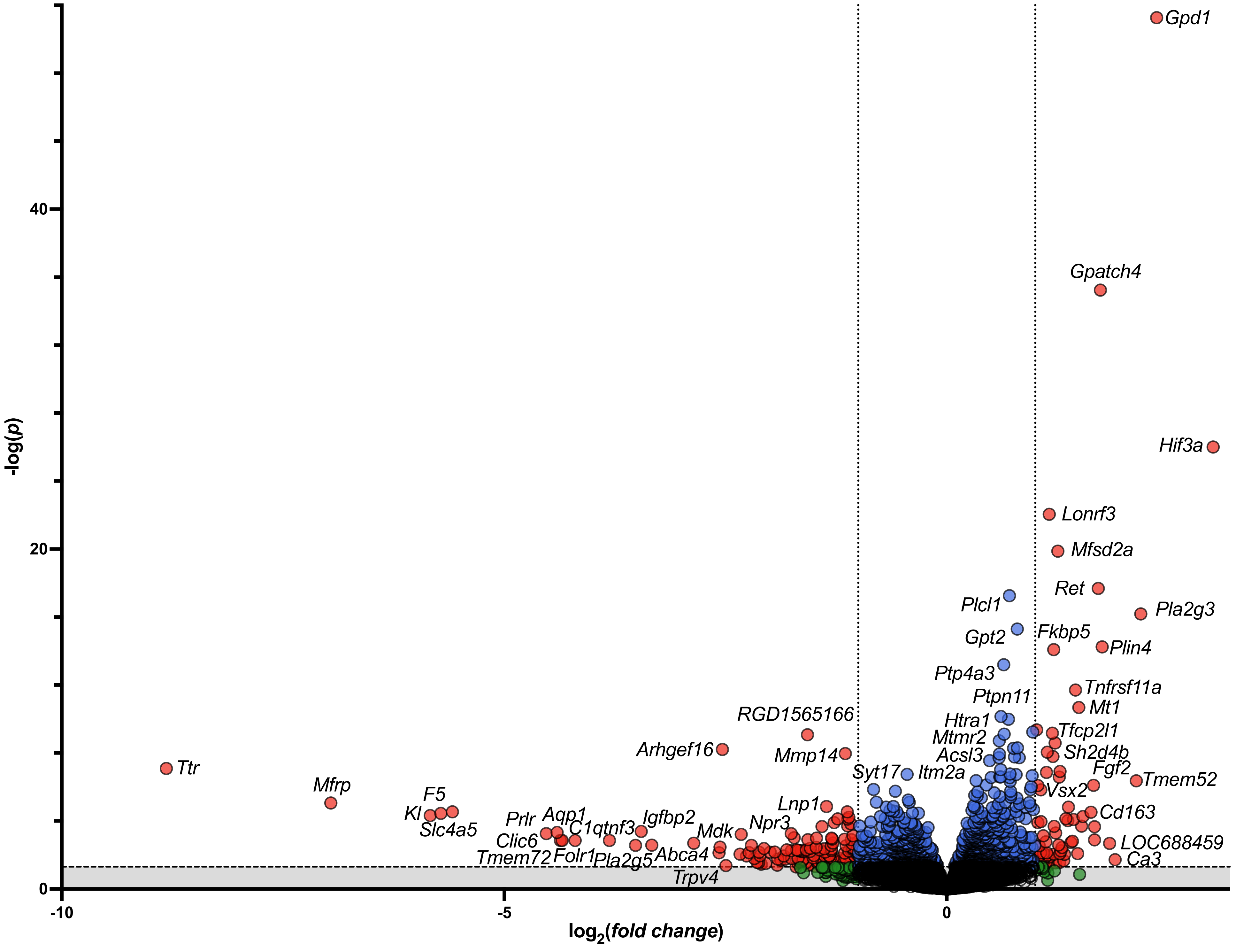

#### *S3. B.* Right Hippocampus 24 Hours Post-SPS

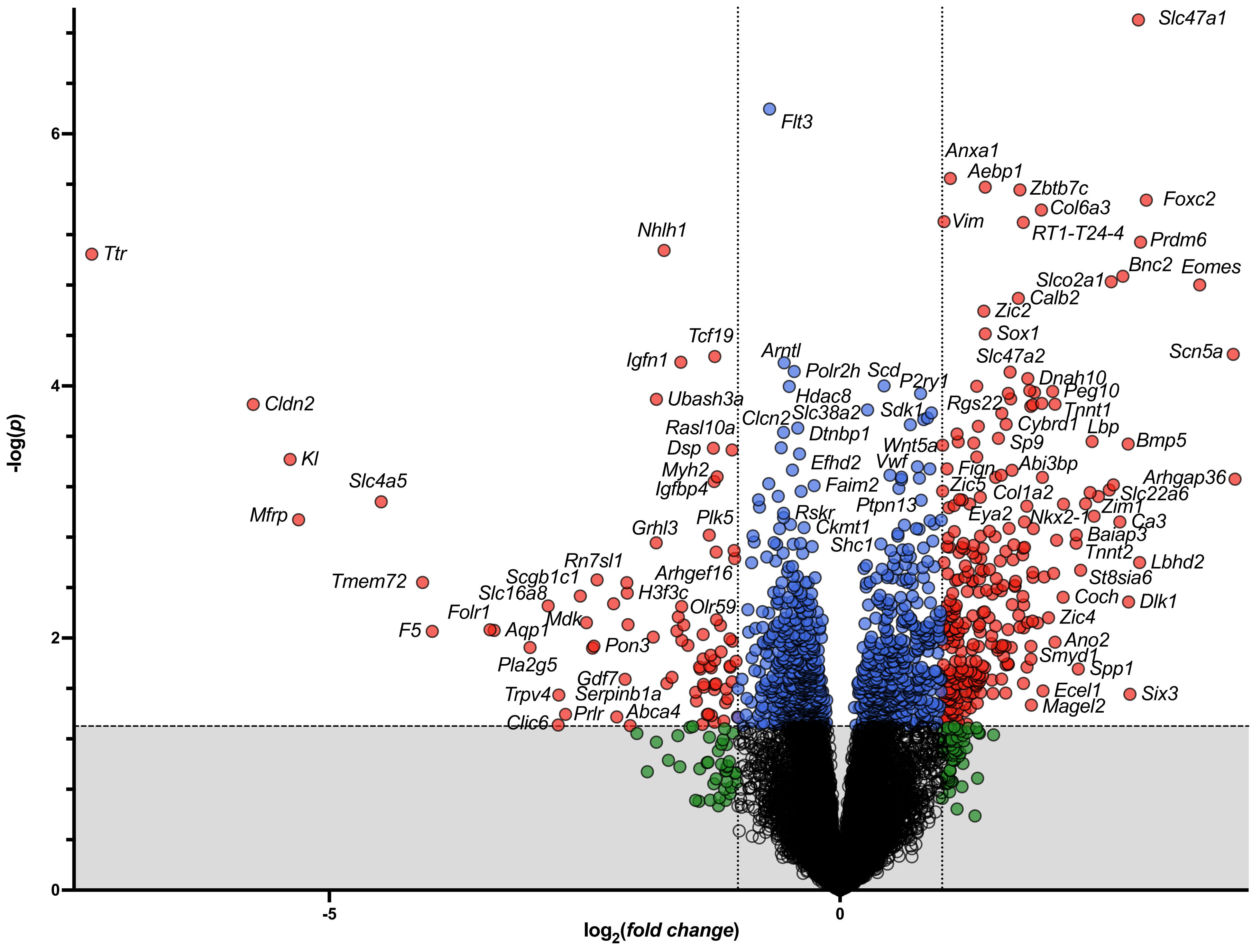

***S3. C.* Right Hippocampus 72 Hours Post-SPS**

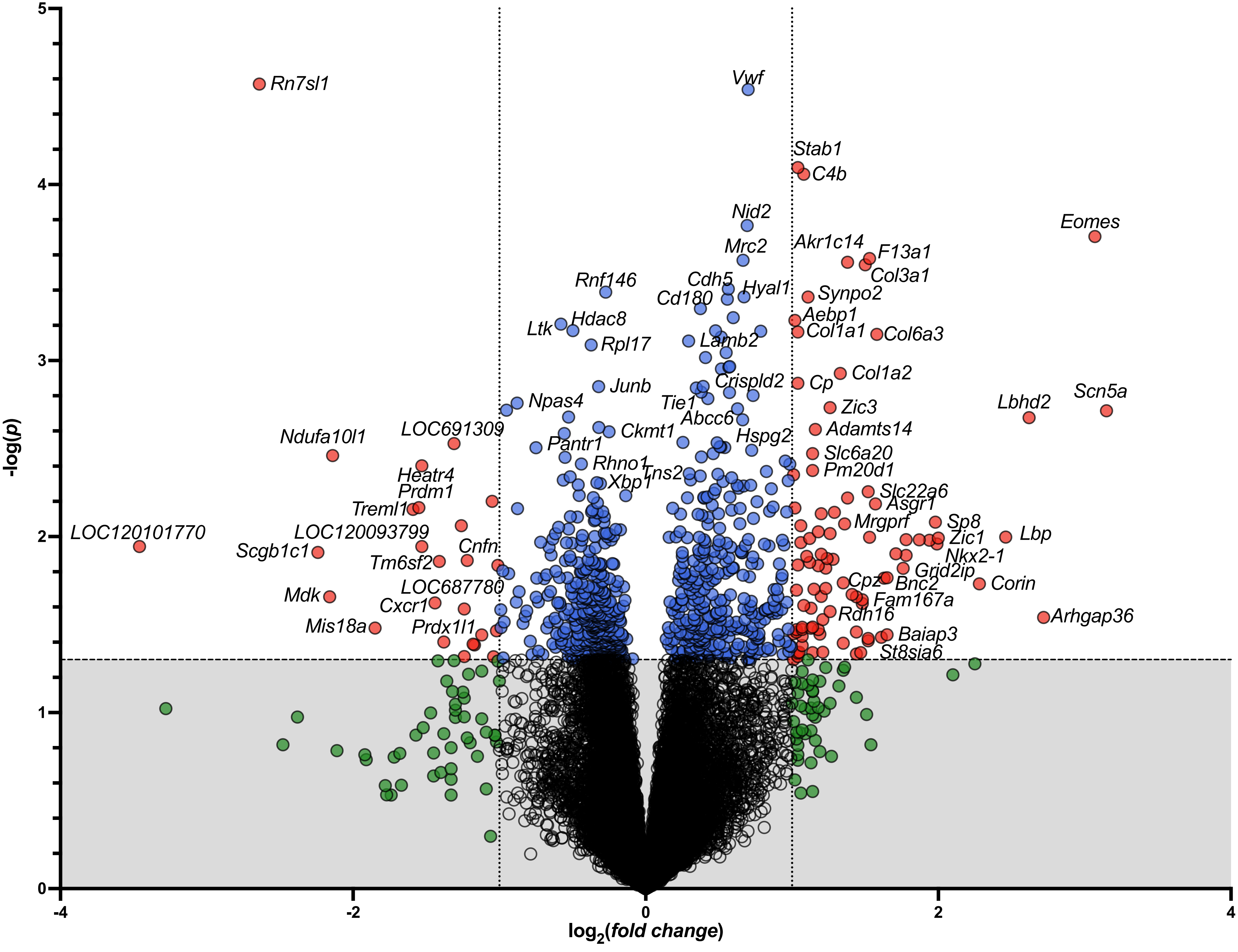

***S3. D.* Left Hippocampus 2 Hours Post-SPS**

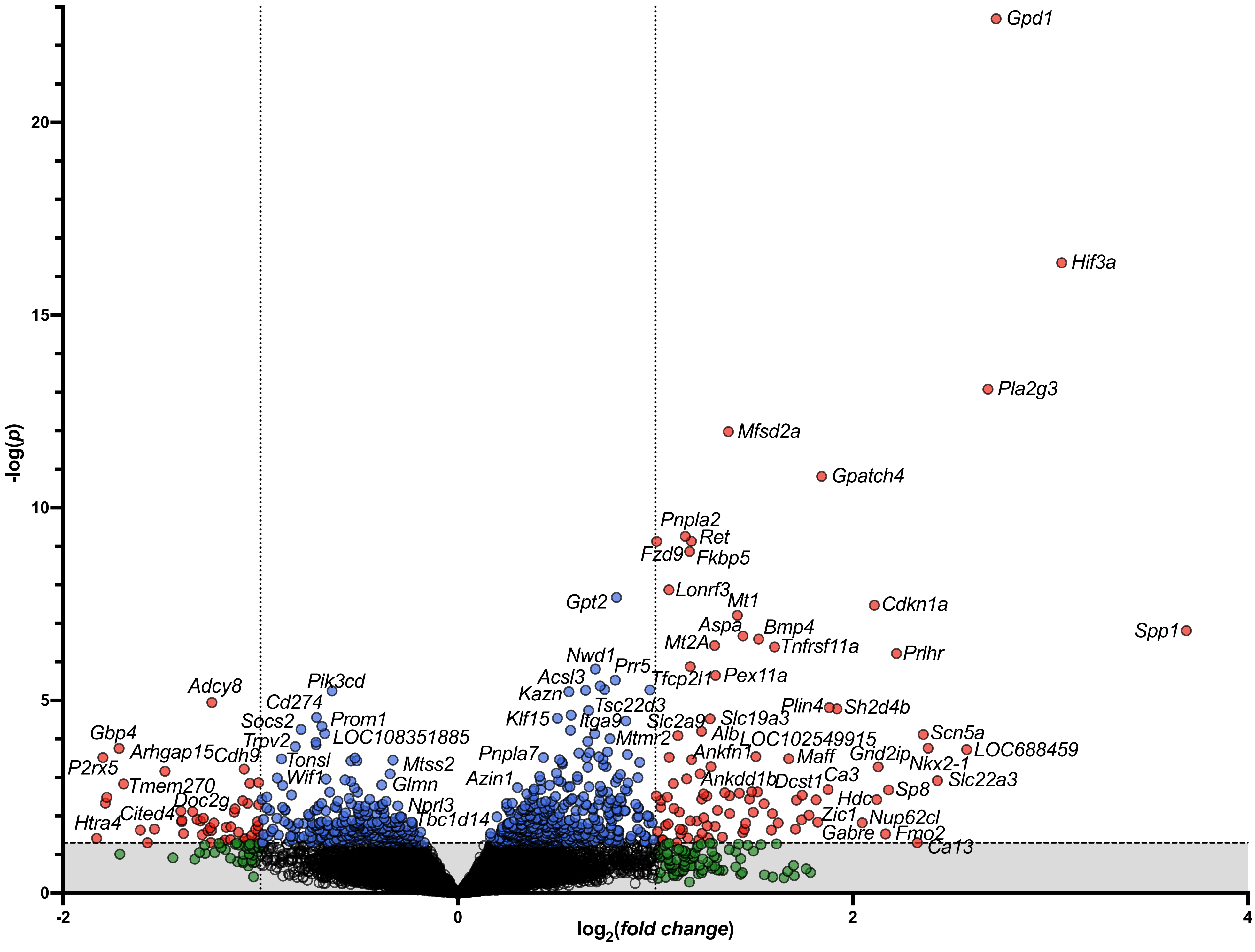

#### *S3. E.* Left Hippocampus 24 Hours Post-SPS

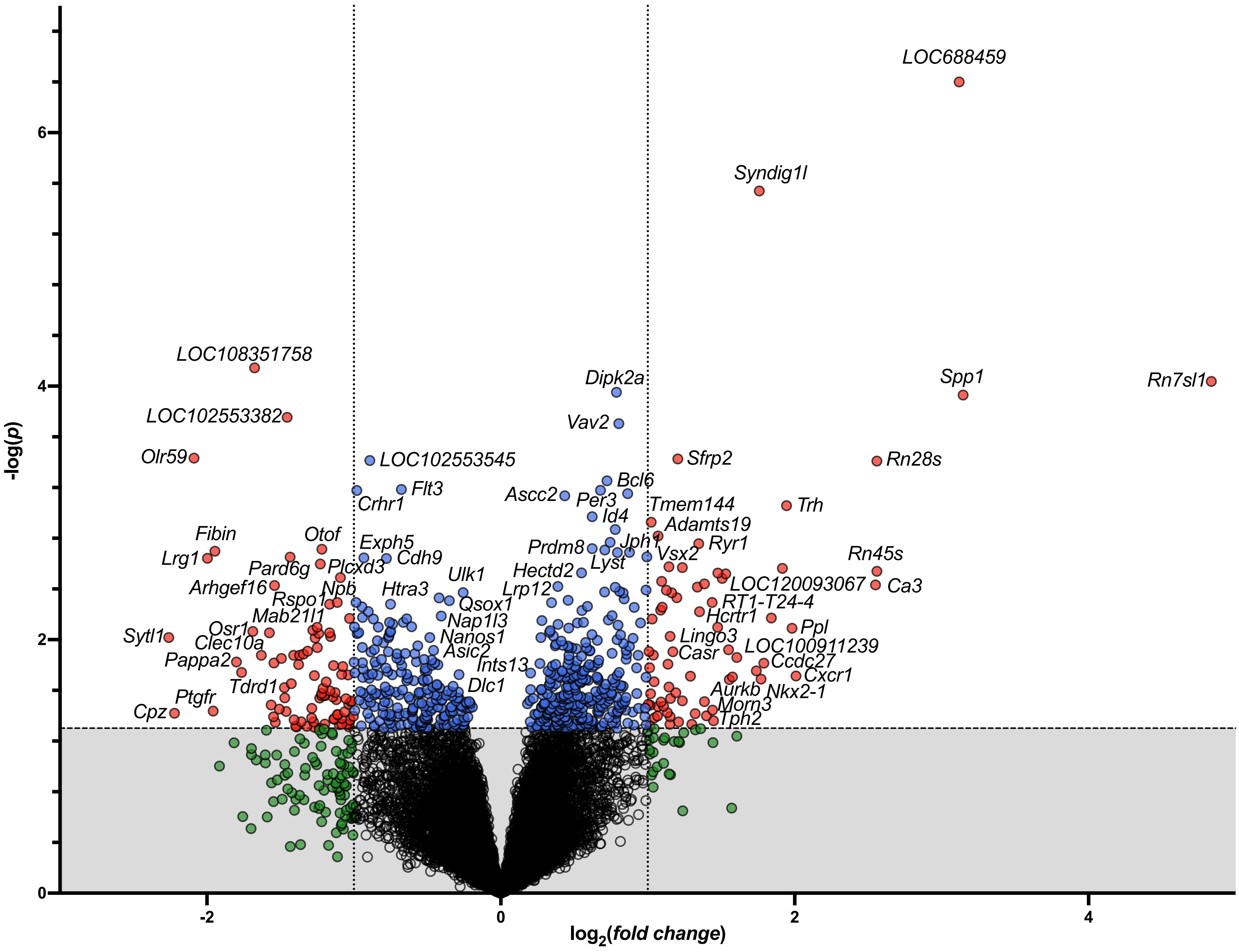

#### *S3. F.* Left Hippocampus 72 Hours Post-SPS

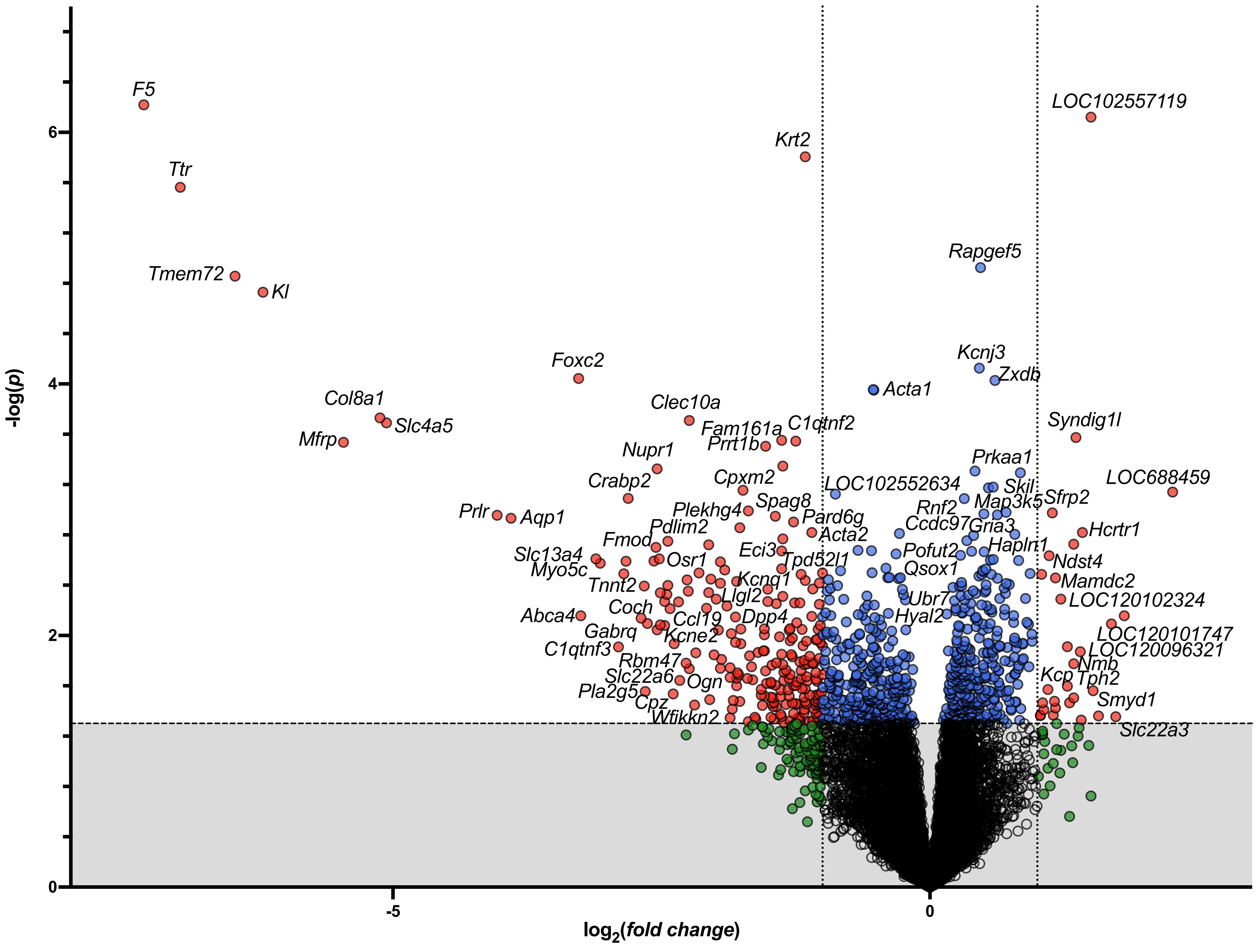

#### *S3. G.* Amygdala 2 Hours Post-SPS

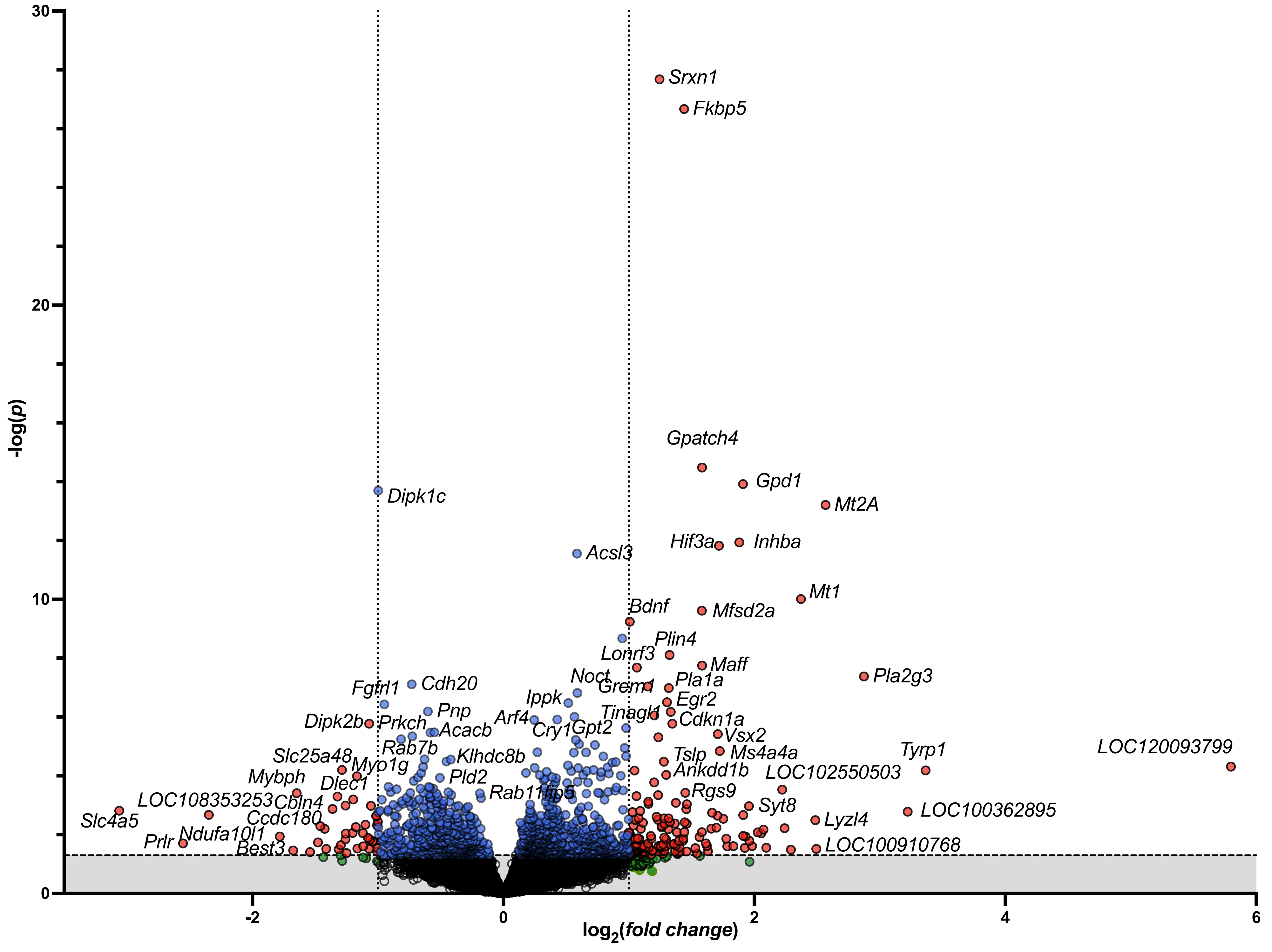

**S3.** ***H.*** **Amygdala 24 Hours Post-SPS**

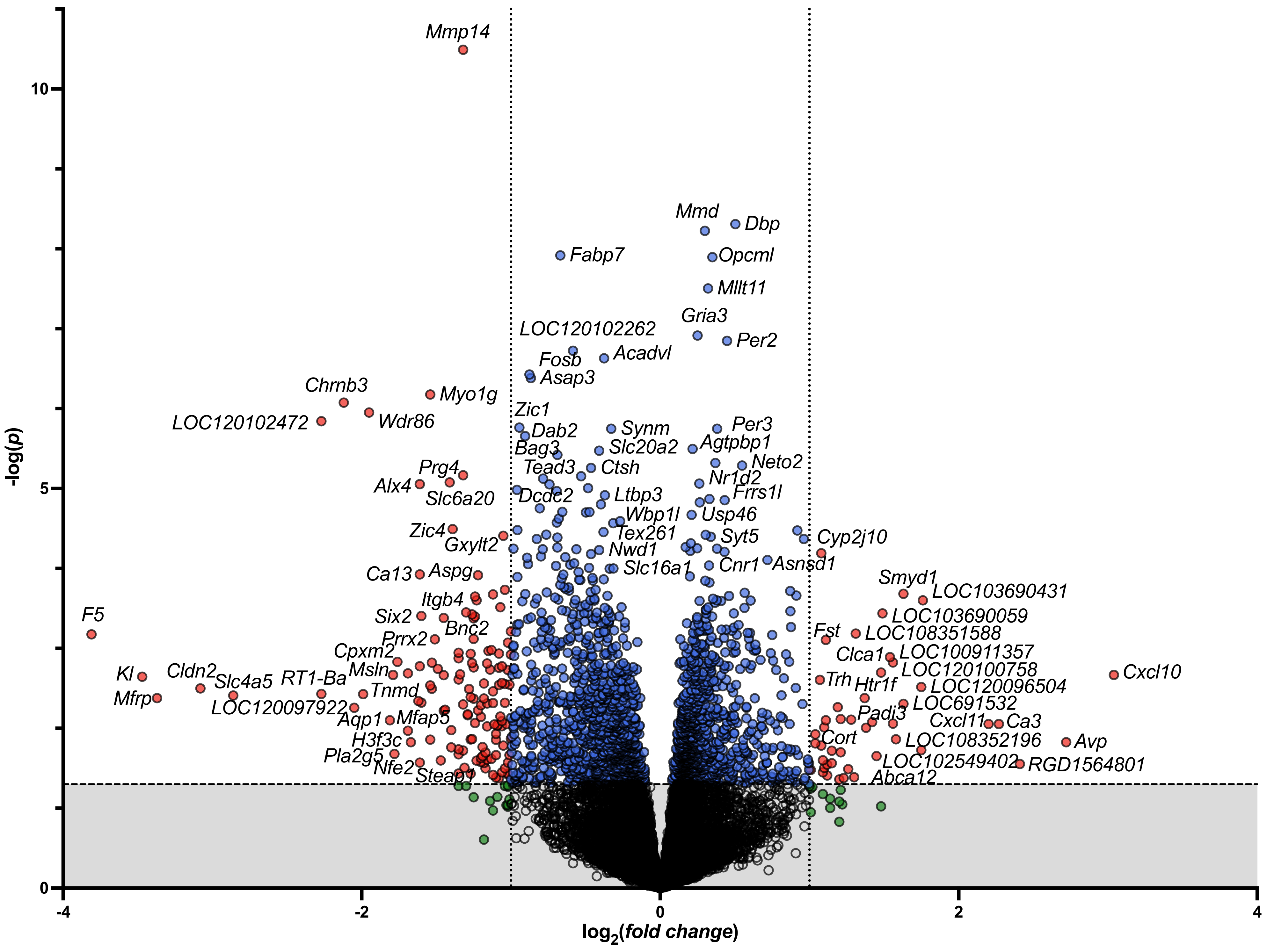

#### *S3. I.* Amygdala 72 Hours Post-SPS

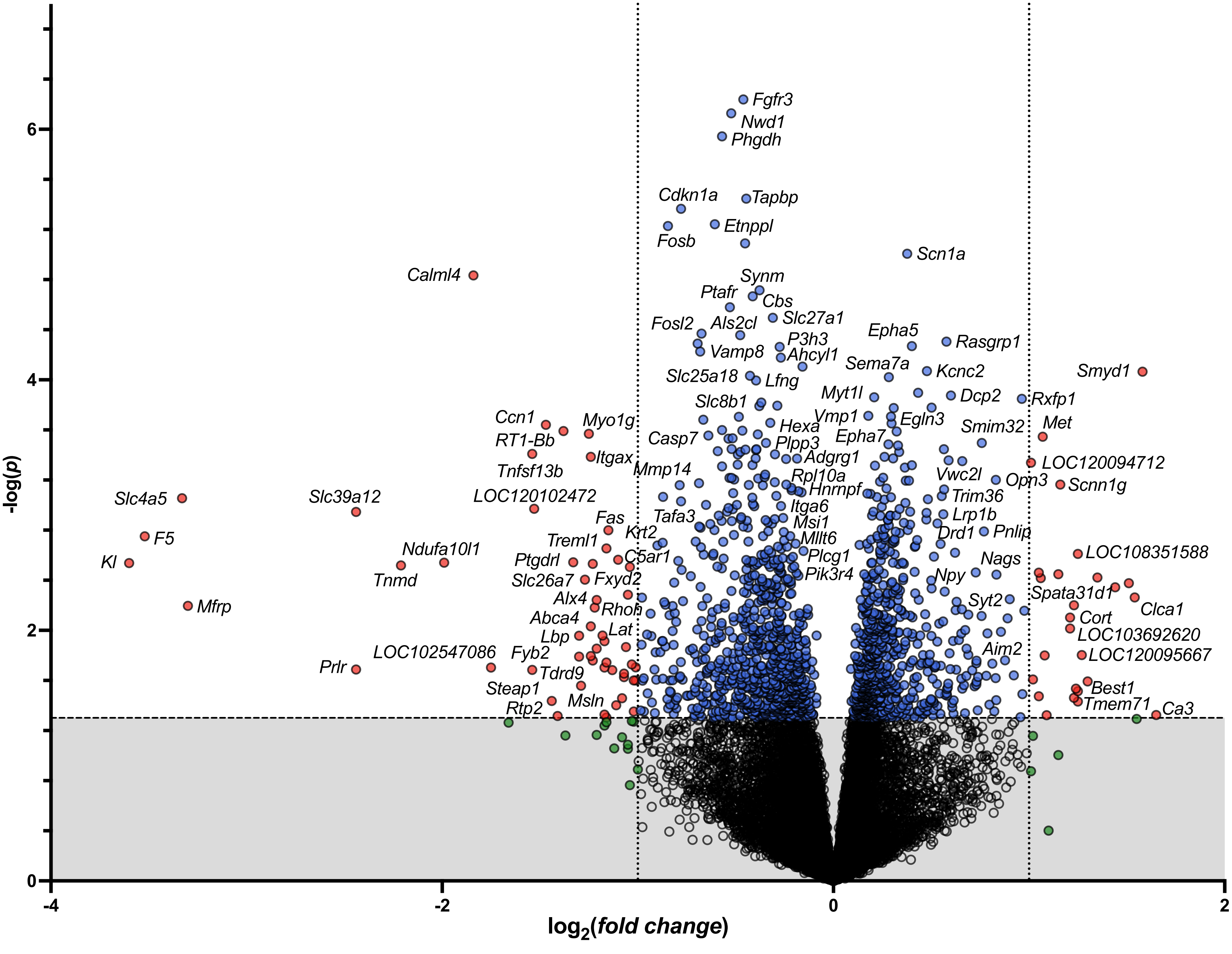

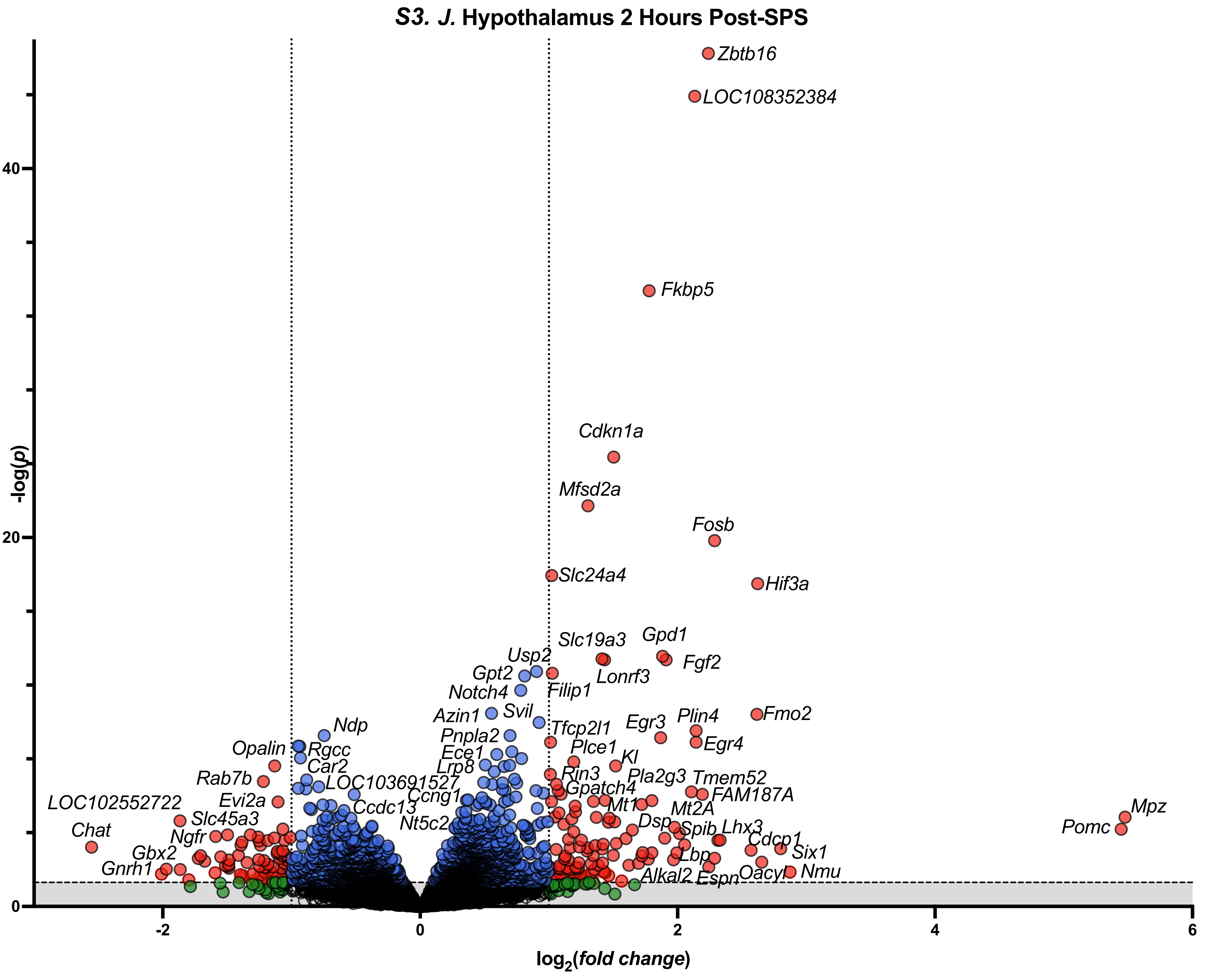

S3. J. Hypothalamus 2 Hours Post-SPS
-log(p)
log2(fold change)
40
20
0
-2
0
2
4
6
Zbtb16
LOC108352384
Fkbp5
Cdkn1a
Mfsd2a
Fosb
Slc24a4
Hif3a
Slc19a3
Gpd1
Usp2
Fgf2
Gpt2
Lonrf3
Notch4
Filip1
Azin1
Svil
Tfcp2l1
Egr3
Plin4
Fmo2
Ndp
Pnpla2
Egr4
Opalin
Rgcc
Ece1
Plce1
Kl
Car2
Lrp8
Rin3
Pla2g3
Tmem52
Rab7b
LOC103691527
Gpatch4
FAM187A
LOC102552722
Evi2a
Ccng1
Mt1
Mt2A
Mpz
Ccdc13
Slc45a3
Nt5c2
Dsp
Spib
Lhx3
Pomc
Chat
Ngfr
Cdcp1
Gbx2
Lbp
Six1
Gnrh1
Oacyl
Nmu
Alkal2
Espn

#### *S3. K.* Hypothalamus 24 Hours Post-SPS

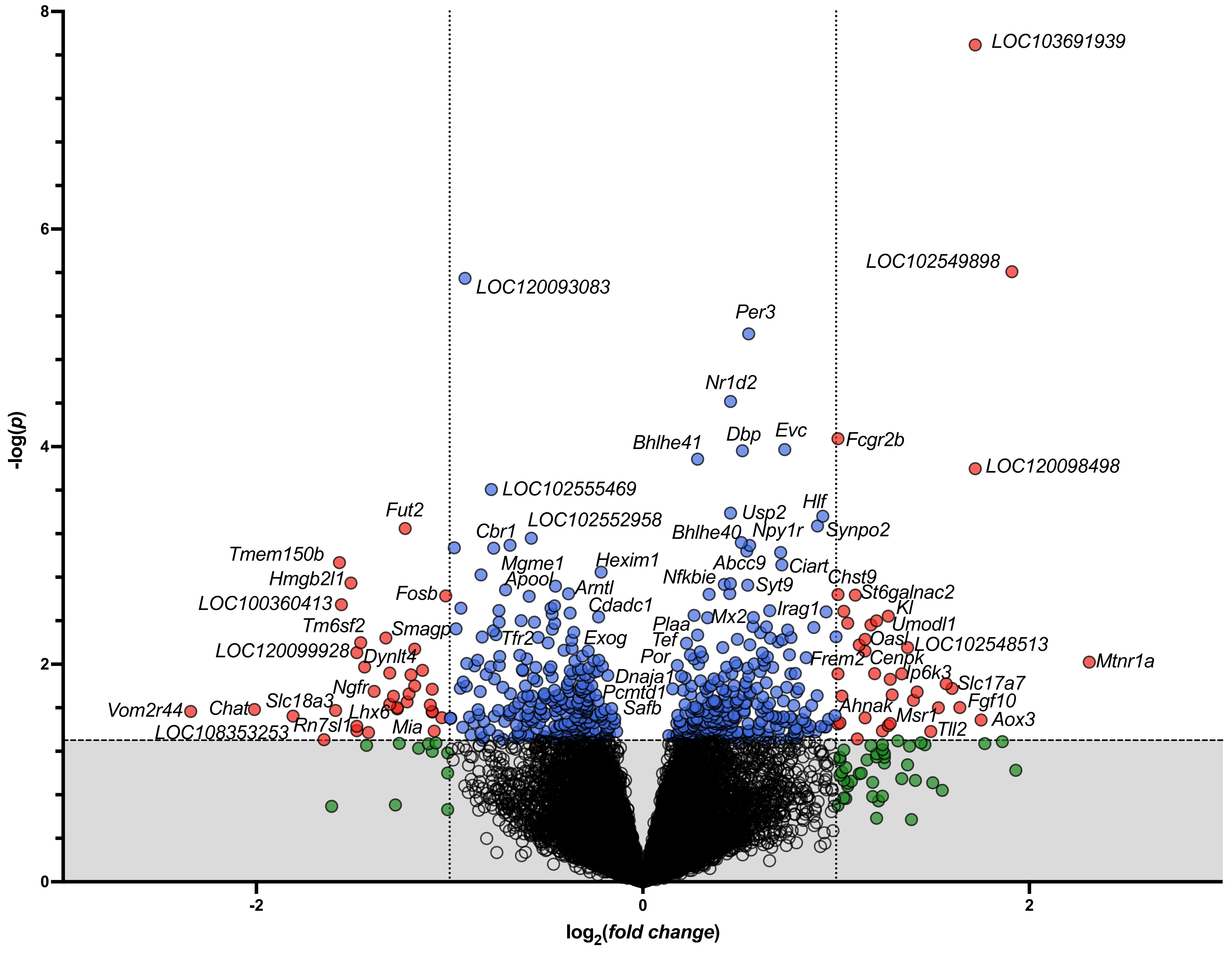

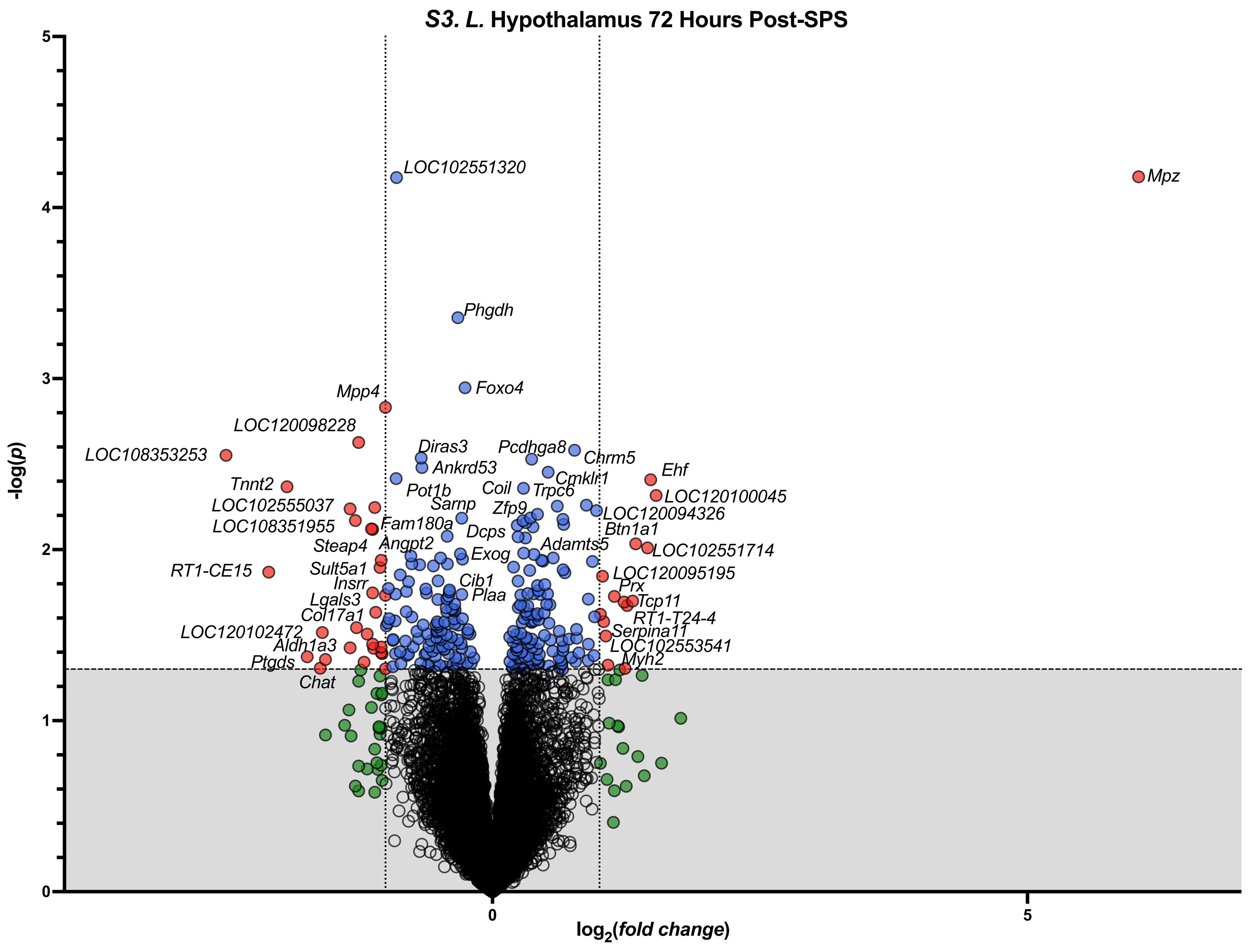

S3. L. Hypothalamus 72 Hours Post-SPS
-log(p)
log2(fold change)
0
1
2
3
4
5
0
5
LOC102551320
Mpz
Phgdh
Foxo4
Mpp4
LOC120098228
LOC108353253
Diras3
Pcdhga8
Chrm5
Ankrd53
Ehf
Cmklr1
Tnnt2
Pot1b
Coil
Trpc6
LOC120100045
LOC102555037
Sarnp
Zfp9
LOC120094326
LOC108351955
Fam180a
Dcps
Btn1a1
Steap4
Angpt2
Adamts5
LOC102551714
Exog
RT1-CE15
Sult5a1
LOC120095195
Cib1
Insrr
Prx
Plaa
Lgals3
Tcp11
Col17a1
RT1-T24-4
LOC120102472
Serpina11
Aldh1a3
LOC102553541
Ptgds
Myh2
Chat

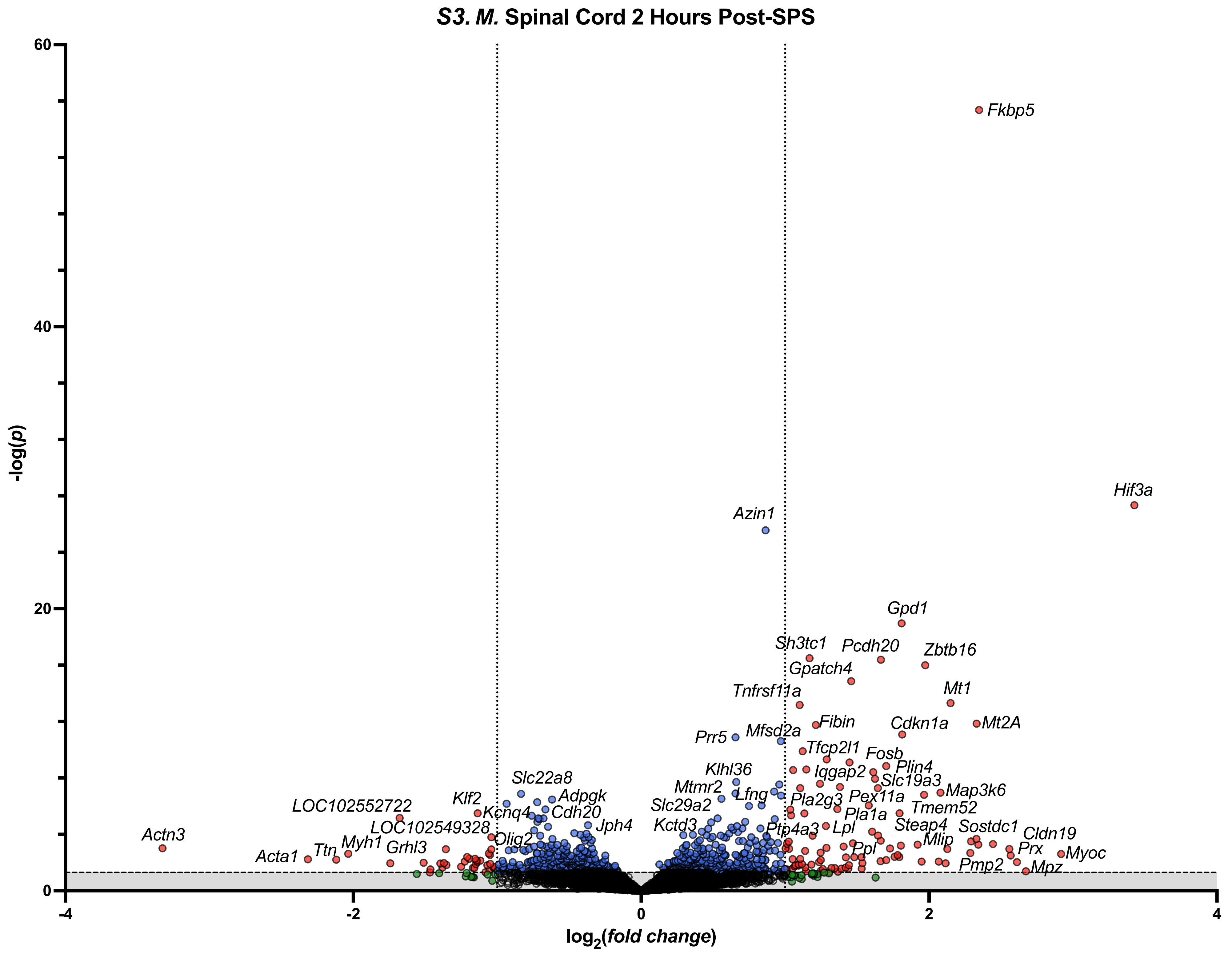

S3. M. Spinal Cord 2 Hours Post-SPS
-log(p)
log2(fold change)
60
40
20
0
-4
-2
0
2
4
Fkbp5
Hif3a
Azin1
Gpd1
Sh3tc1
Pcdh20
Zbtb16
Gpatch4
Mt1
Tnfrsf11a
Fibin
Cdkn1a
Mt2A
Prr5
Mfsd2a
Tfcp2l1
Fosb
Klhl36
Iqgap2
Plin4
Slc19a3
Mtmr2
Map3k6
Lfng
Pla2g3
Pex11a
Slc29a2
Tmem52
Pla1a
Kctd3
Ptp4a3
Lpl
Steap4
Sostdc1
Cldn19
Mlip
Ppl
Prx
Myoc
Pmp2
Mpz
Slc22a8
Adpgk
Cdh20
Klf2
Kcnq4
LOC102552722
LOC102549328
Jph4
Olig2
Actn3
Myh1
Grhl3
Ttn
Acta1

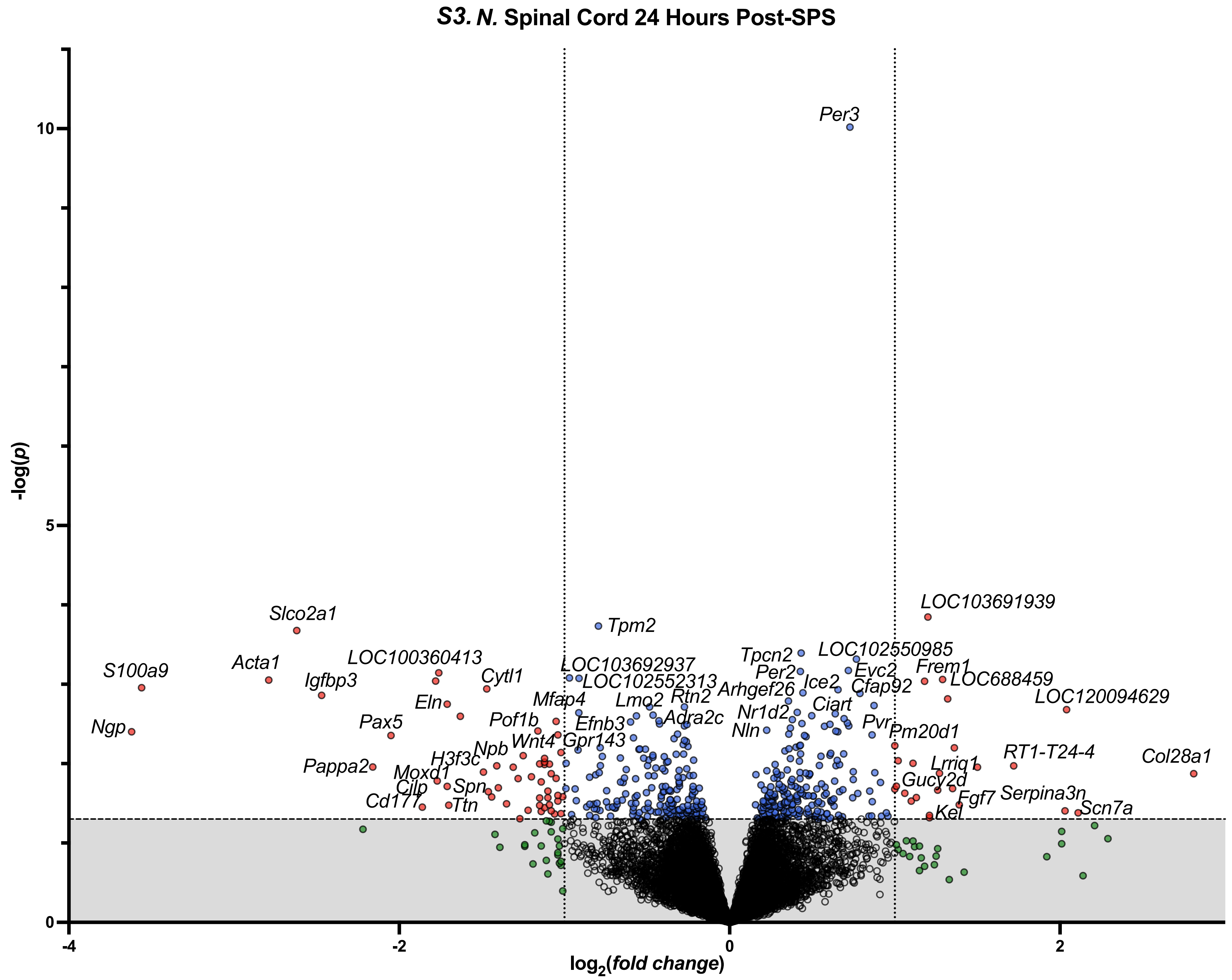

S3. N. Spinal Cord 24 Hours Post-SPS
-log(p)
log2(fold change)
10
5
0
-4
-2
0
2
Per3
Tpm2
Slco2a1
S100a9
Acta1
Igfbp3
Ngp
LOC100360413
Cytl1
Eln
Pax5
Pof1b
Mfap4
LOC103692937
LOC102552313
Lmo2
Rtn2
Efnb3
Adra2c
Wnt4
Gpr143
Npb
H3f3c
Pappa2
Moxd1
Cilp
Spn
Cd177
Ttn
Tpcn2
Per2
Arhgef26
Ice2
Nr1d2
Nln
Ciart
LOC102550985
Evc2
Cfap92
Pvr
Pm20d1
LOC103691939
Frem1
LOC688459
LOC120094629
RT1-T24-4
Col28a1
Lrriq1
Gucy2d
Fgf7
Serpina3n
Scn7a
Kel

**S3. O. Spinal Cord 72 Hours Post-SPS**

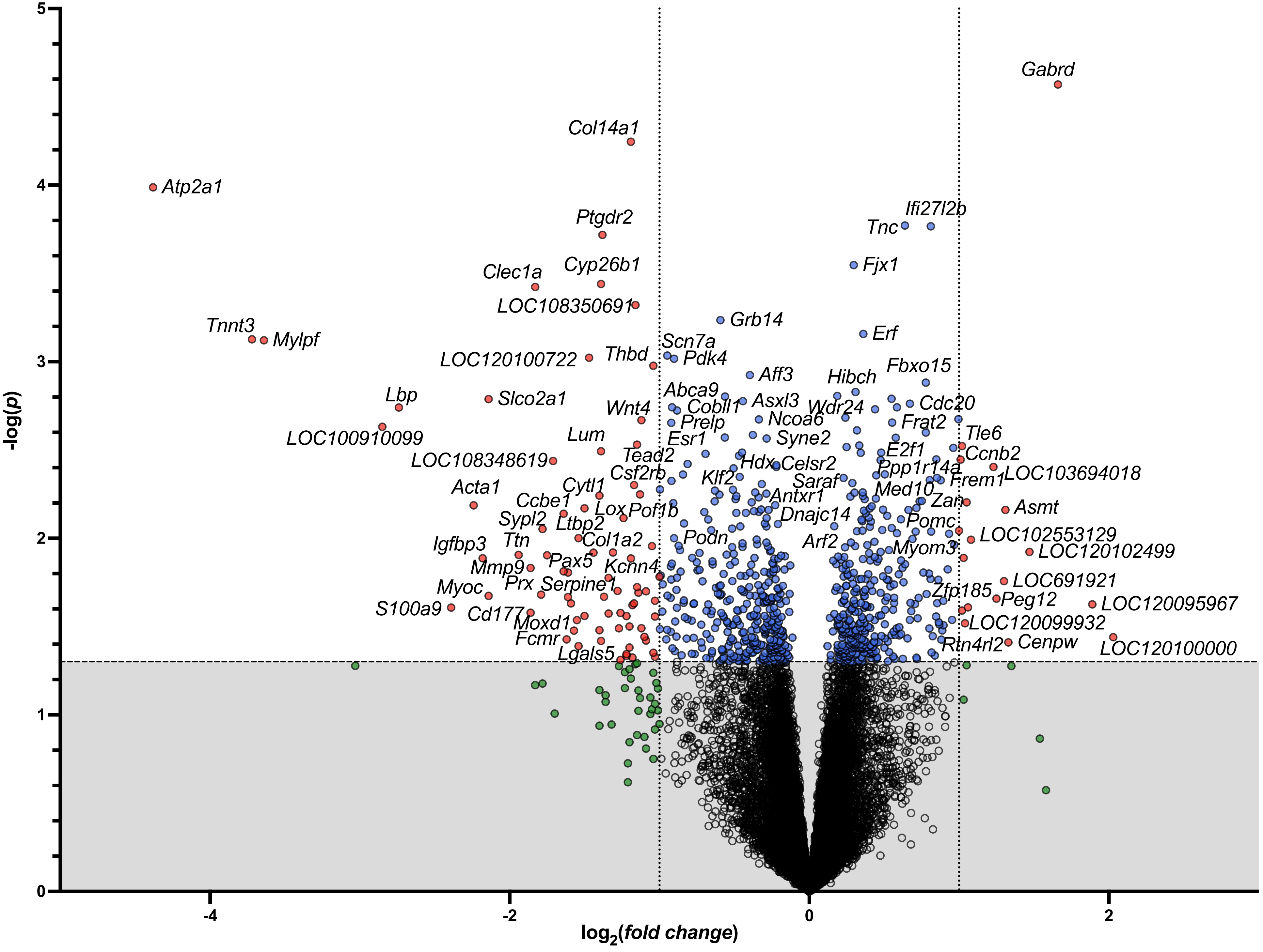

**S3. P. Dorsal Root Ganglia 2 Hours Post-SPS**

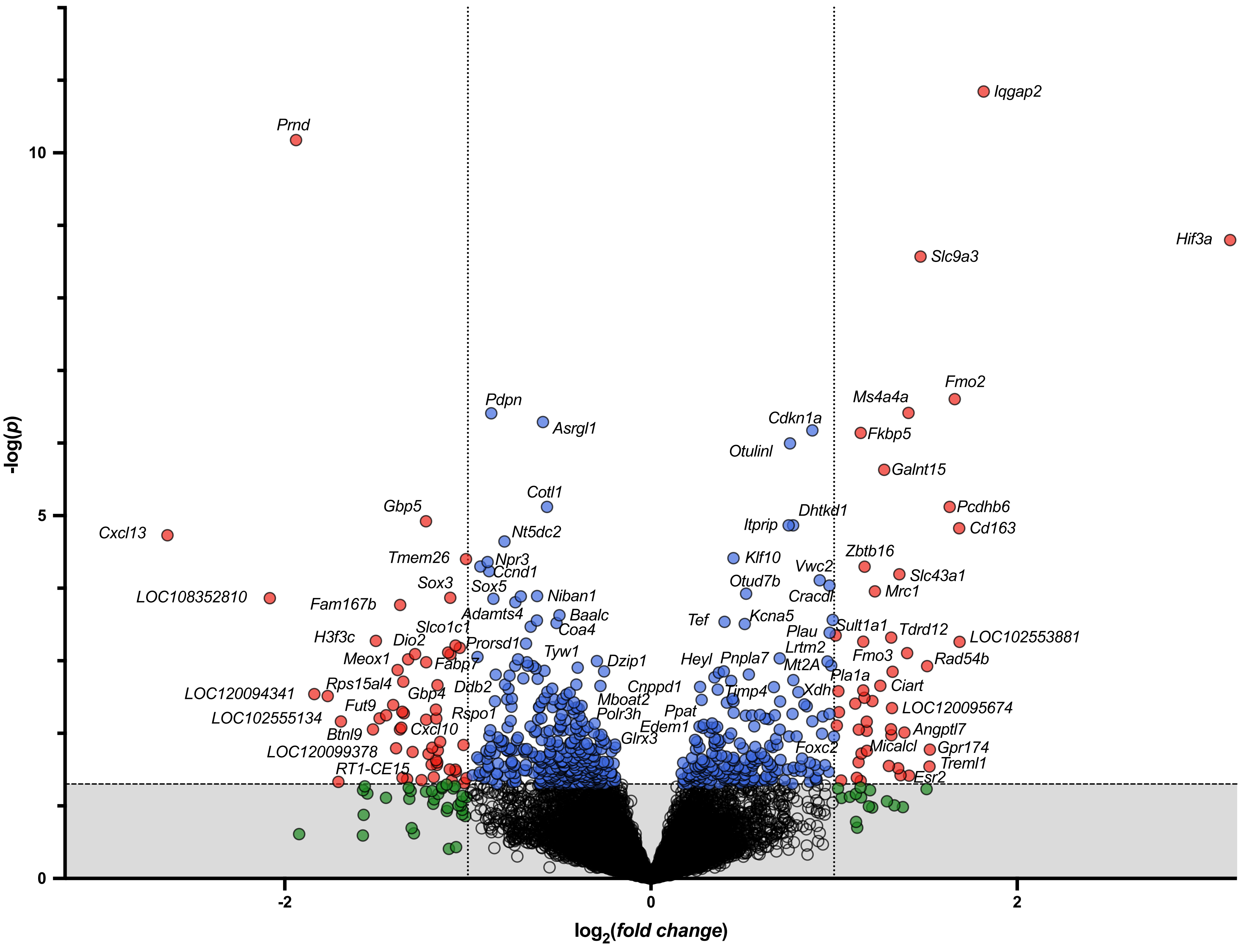

***S3. Q.*** **Dorsal Root Ganglia 24 Hours Post-SPS**

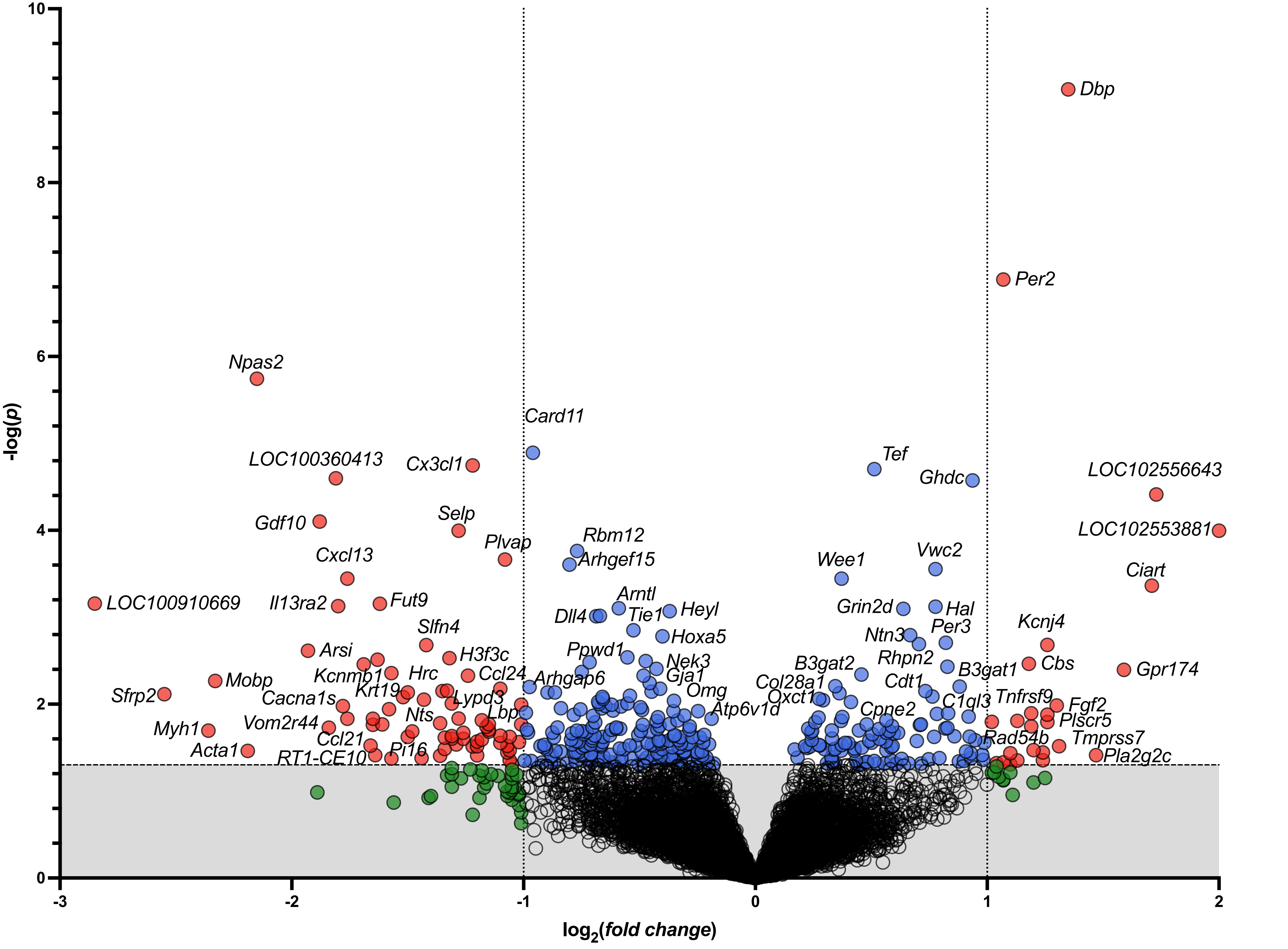

***S3. R.* Dorsal Root Ganglia 72 Hours Post-SPS**

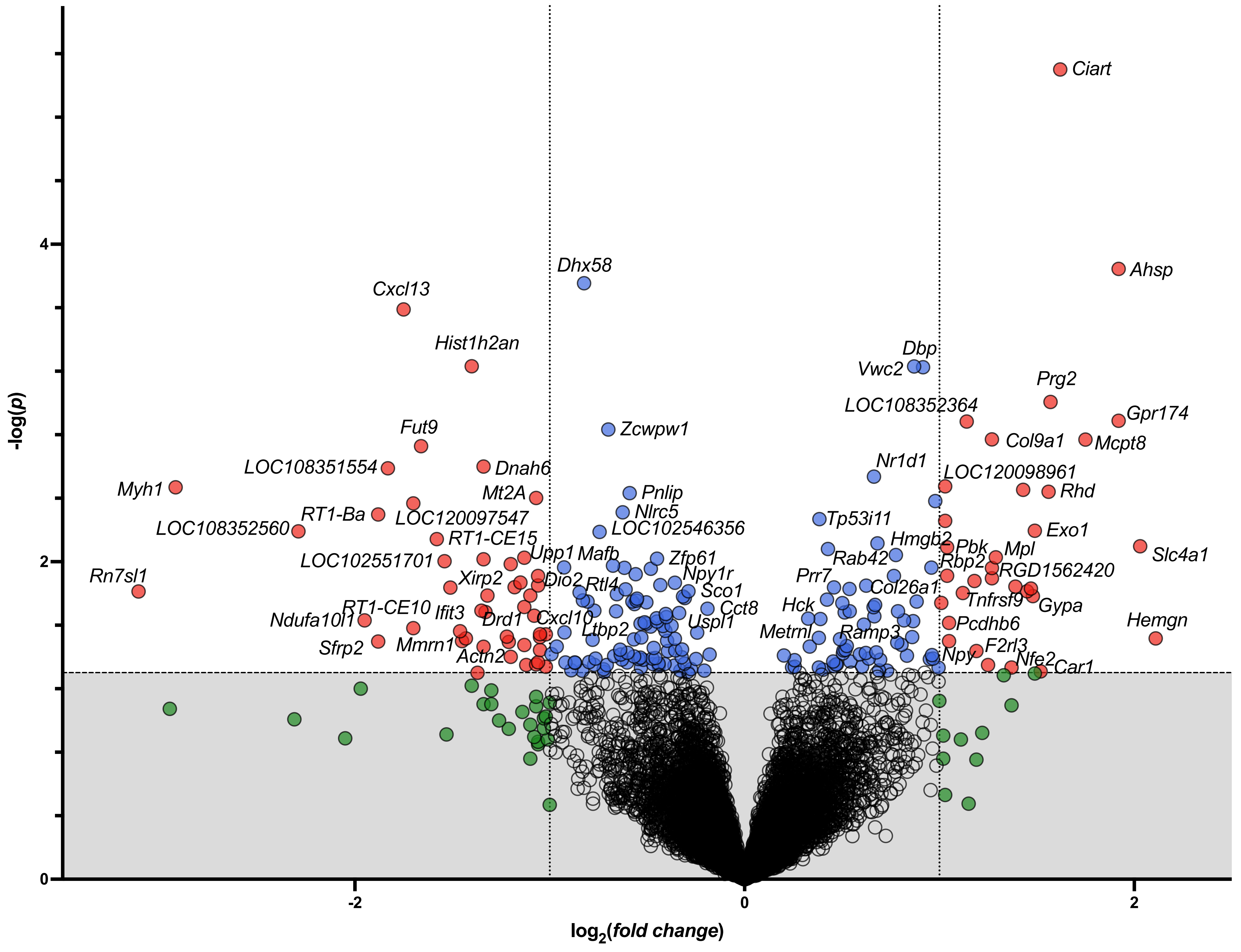

**S3. S. Muscle 2 Hours Post-SPS**

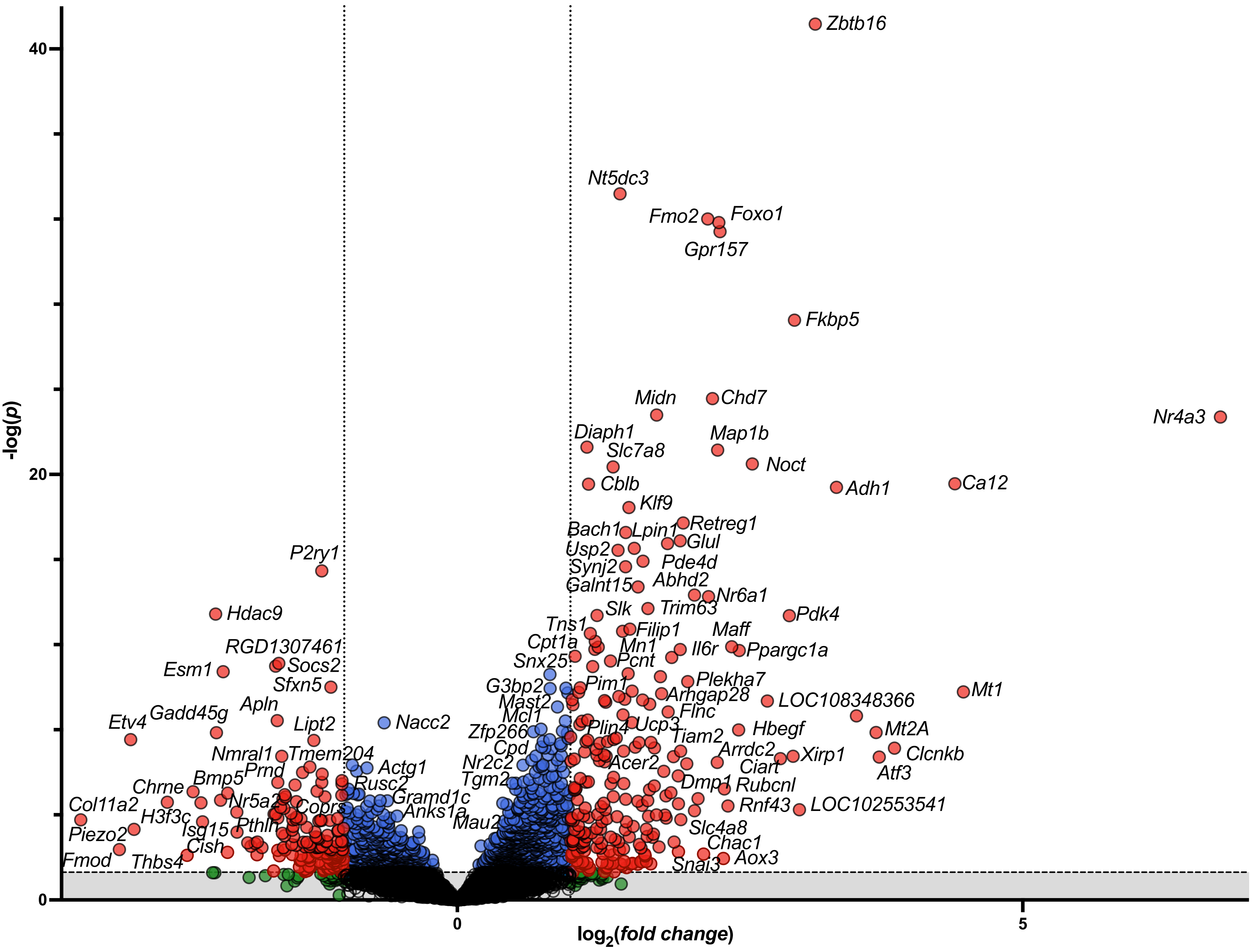

#### *S3. T.* Muscle 24 Hours Post-SPS

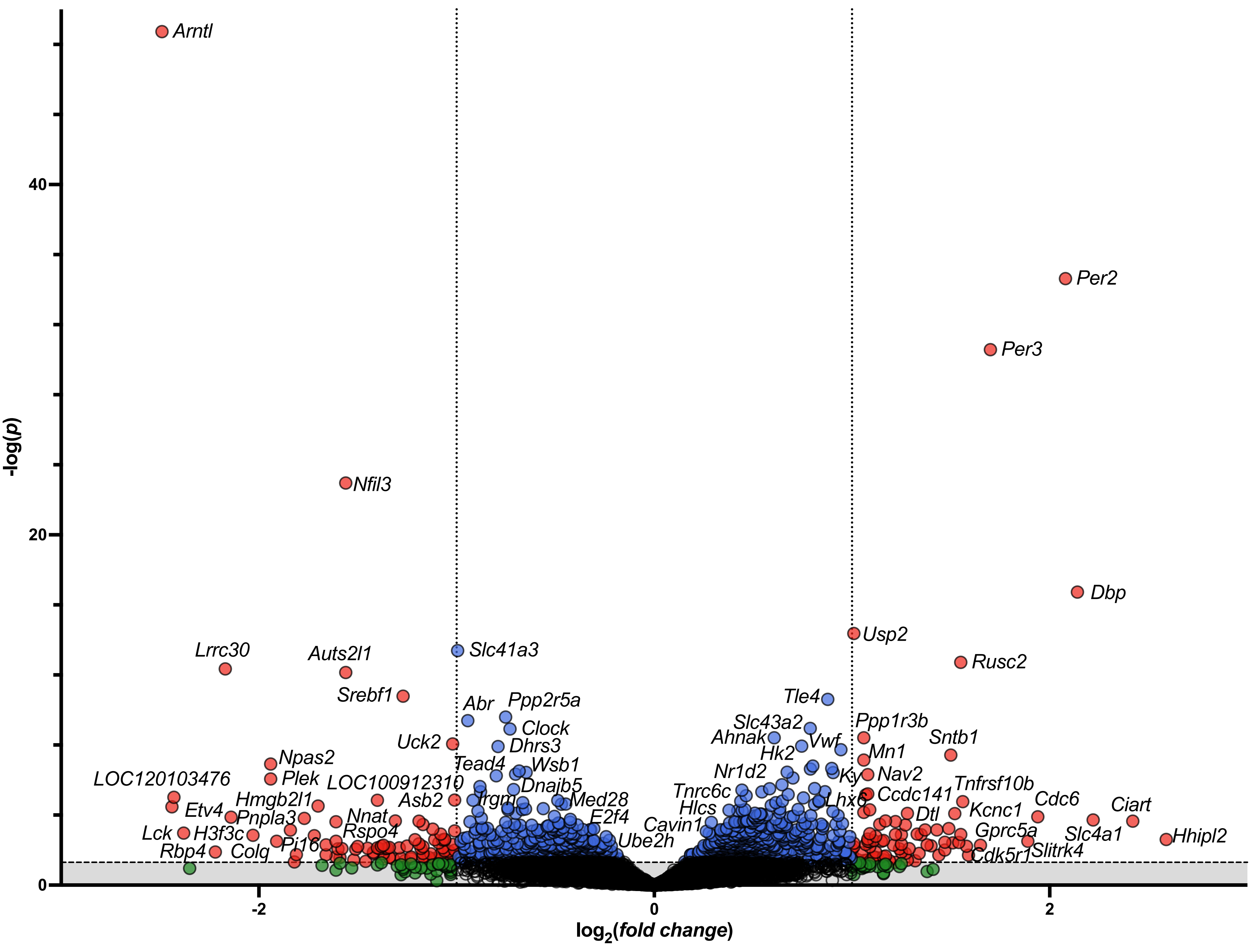

**S3. U. Muscle 72 Hours Post-SPS**

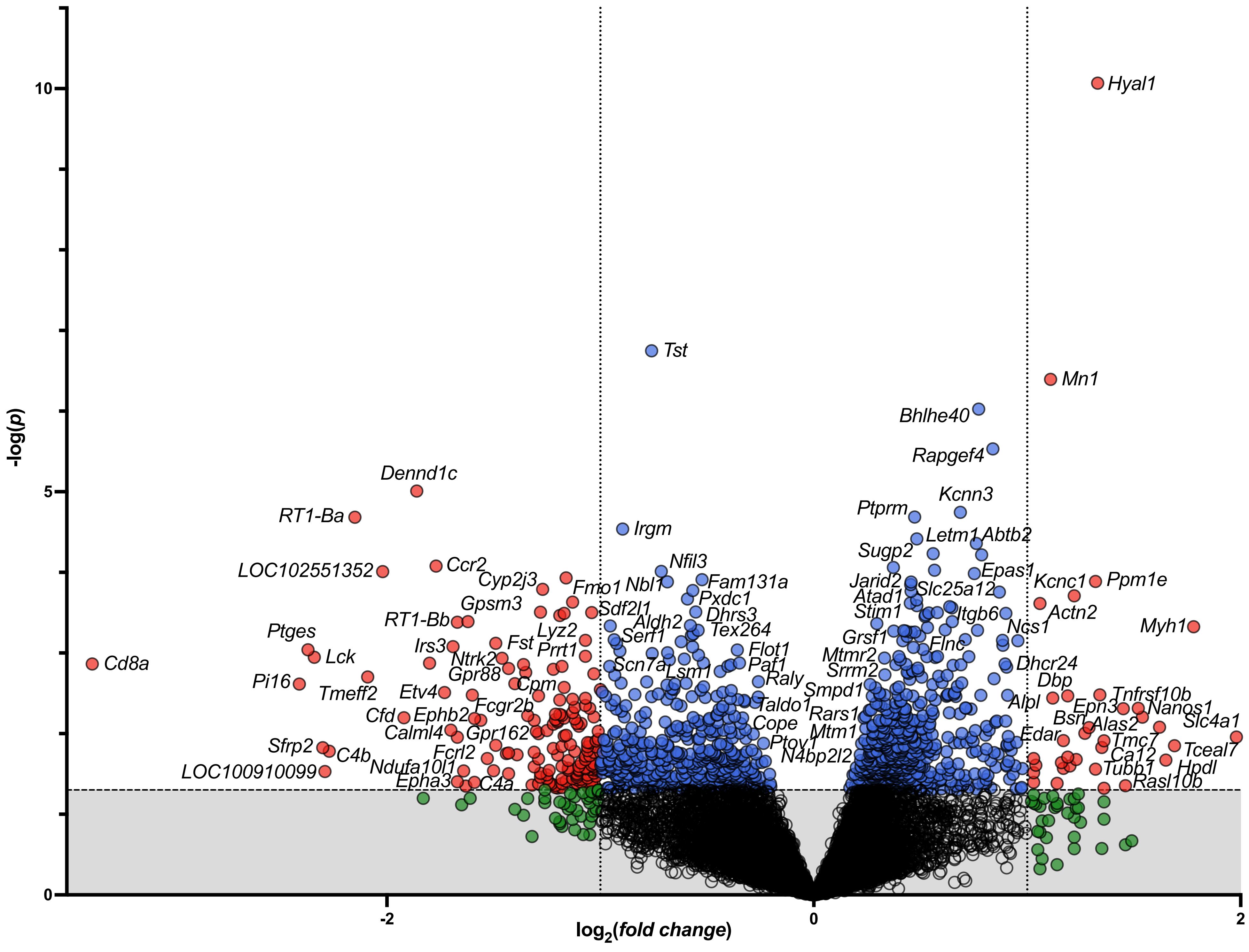

***S3. V.* Heart 2 Hours Post-SPS**

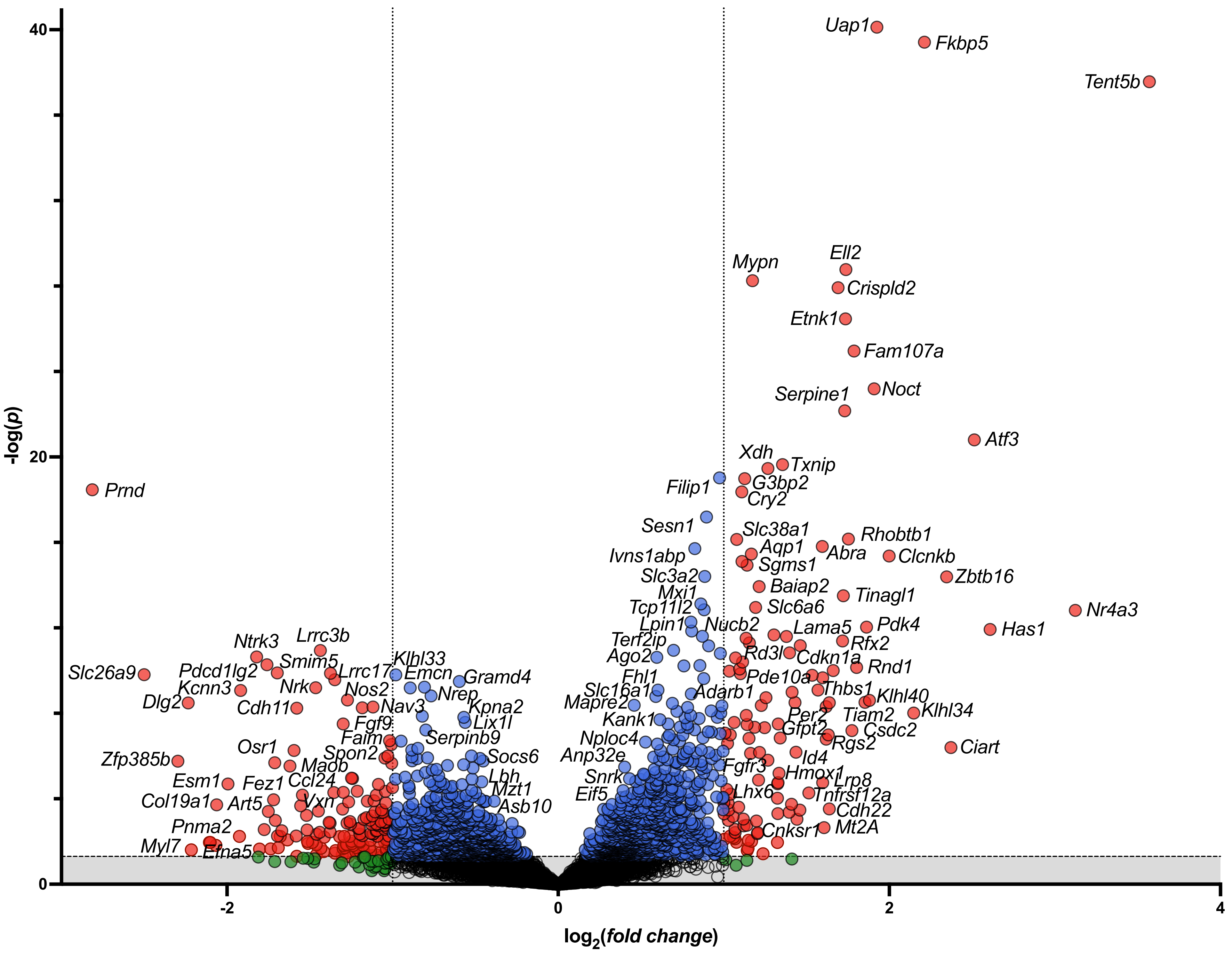

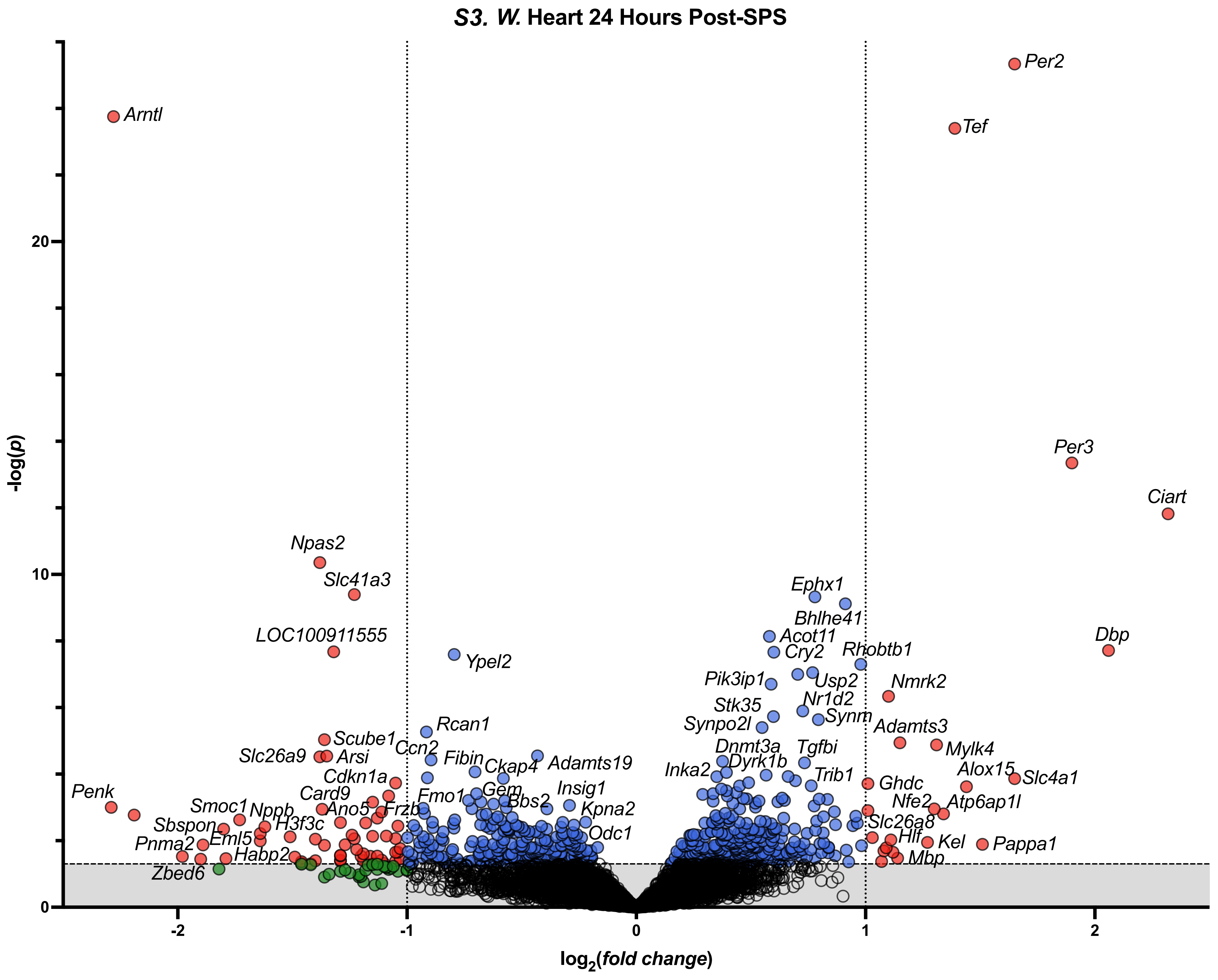

S3. W. Heart 24 Hours Post-SPS
-log(p)
log2(fold change)
0
10
20
-2
-1
0
1
2
Arntl
Per2
Tef
Per3
Ciart
Dbp
Npas2
Slc41a3
LOC100911555
Ypel2
Ephx1
Bhlhe41
Acot11
Cry2
Rhobtb1
Pik3ip1
Usp2
Nmrk2
Nr1d2
Stk35
Synm
Synpo2l
Adamts3
Rcan1
Scube1
Ccn2
Slc26a9
Arsi
Fibin
Ckap4
Adamts19
Dnmt3a
Tgfbi
Mylk4
Dyrk1b
Inka2
Alox15
Slc4a1
Trib1
Cdkn1a
Ghdc
Penk
Card9
Fmo1
Gem
Insig1
Bbs2
Kpna2
Nfe2
Atp6ap1l
Smoc1
Nppb
Ano5
Frzb
Slc26a8
Sbspon
H3f3c
Odc1
Hlf
Kel
Pappa1
Pnma2
Eml5
Habp2
Mbp
Zbed6

S3. X. Heart 72 Hours Post-SPS

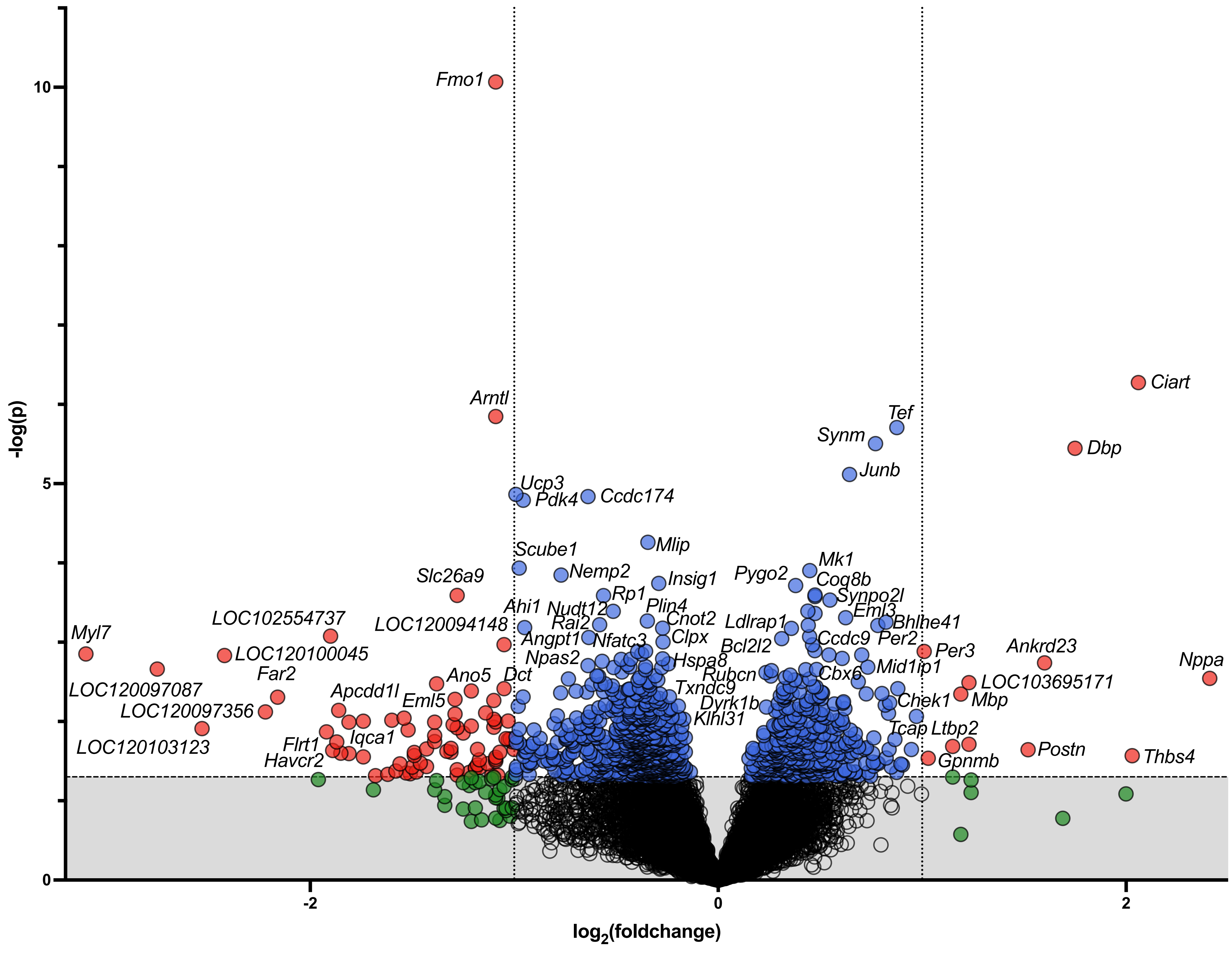

| | 2 hours | | | | 24 hours | | | | 72 hours | | | |
|---|---|---|---|---|---|---|---|---|---|---|---|---|
| **Right Hippocampus** | ↓↓↓ | Ttr | ↑ | Fkbp5 | ↓↓↓ | Ttr | ↑↑ | Bnc2 | ↓↓ | Rn7sl1 | ↑ | Akr1c14 |
| | ↓↓↓ | Mfrp | ↑ | Mfsd2a | ↓↓↓ | Cldn2 | ↑↑ | Slco2a1 | ↑↑ | Eomes | ↑ | C4b |
| | ↓↓↓ | Kl | ↑ | Tnfrsf11a | ↑↑ | Slc47a1 | ↑↑ | Arhgap36 | ↑↑ | Scn5a | ↓ | Scgb1c1 |
| | ↓↓↓ | F5 | ↑ | Mt1 | ↓↓↓ | Kl | ↑ | Col6a3 | ↑↑ | Lbhd2 | ↑ | Stab1 |
| | ↓↓↓ | Slc4a5 | ↑ | Ret | ↑↑ | Eomes | ↑↑ | Bmp5 | ↓↓ | LOC120101770 | ↑↑ | Arhgap36 |
| | ↓↓ | Prlr | ↑ | Gpatch4 | ↑↑ | Foxc2 | ↓↓ | Tmem72 | ↑ | F13a1 | ↑ | Sp8 |
| | ↓↓ | Aqp1 | ↑ | Plin4 | ↑↑ | Scn5a | ↑ | Zbtb7c | ↑ | Col3a1 | ↑ | Zic1 |
| | ↓↓ | Arhgef16 | ↑ | Pla2g3 | ↓↓↓ | Mfrp | ↑ | RT1-T24-4 | ↓ | Ndufa10l1 | ↑ | Corin |
| | ↓ | RGD1565166 | ↑ | Gpd1 | ↑↑ | Prdm6 | ↑ | Lbp | ↑ | Col6a3 | ↑ | Col1a2 |
| | ↑ | Lonrf3 | ↑↑ | Hif3a | ↓↓ | Slc4a5 | ↓ | Nhlh1 | ↑ | Lbp | ↑ | Nkx2-1 |
| **Left Hippocampus** | ↑↑ | Gpd1 | ↑ | Fkbp5 | ↑↑ | LOC688459 | ↑ | LOC102553545 | ↓↓↓ | F5 | ↑ | Acta1 |
| | ↑↑ | Hif3a | ↑ | Tnfrsf11a | ↑ | Syndig1l | ↑↑ | Rn28s | ↑ | LOC102557119 | ↑ | LOC102548599 |
| | ↑↑ | Pla2g3 | ↑ | Mt1 | ↓ | LOC108351758 | ↑ | Bcl6 | ↓ | Krt2 | ↓↓↓ | Col8a1 |
| | ↑↑ | Spp1 | ↑ | Bmp4 | ↑↑ | Rn7sl1 | ↑ | Flt3 | ↓↓↓ | Ttr | ↓ | Clec10a |
| | ↑ | Gpatch4 | ↑ | Scn5a | ↑ | Dipk2a | ↑ | Per3 | ↑ | Rapgef5 | ↓↓↓ | Slc4a5 |
| | ↑ | Mfsd2a | ↑ | Aspa | ↑↑ | Spp1 | ↑ | Crhr1 | ↓↓↓ | Tmem72 | ↑ | Syndig1l |
| | ↑ | Cdkn1a | ↑↑ | LOC688459 | ↓ | LOC102553382 | ↑ | Ntf3 | ↓↓↓ | Kl | ↓ | Fam161a |
| | ↑ | Prlhr | ↑ | Fzd9 | ↑ | Vav2 | ↑ | Ascc2 | ↑ | Kcnj3 | ↓ | C1qtnf2 |
| | ↑ | Pnpla2 | ↑ | Sh2d4b | ↓ | Olr59 | ↑ | Trh | ↓↓ | Foxc2 | ↓↓↓ | Mfrp |
| | ↑ | Ret | ↑ | Plin4 | ↑ | Sfrp2 | ↑ | Id4 | ↑ | Zxdb | ↓ | Prrt1b |
| **Spinal Cord** | ↑ | Fkbp5 | ↑ | Sh3tc1 | ↓↓ | S100a9 | ↓ | Pax5 | ↓↓ | Atp2a1 | ↓ | Acta1 |
| | ↑↑ | Hif3a | ↑ | Fosb | ↓↓ | Slco2a1 | ↓ | Eln | ↓↓ | Tnnt3 | ↓ | Cyp26b1 |
| | ↑ | Gpd1 | ↑ | Tnfrsf11a | ↓↓ | Ngp | ↑ | LOC103691939 | ↓↓ | Mylpf | ↓ | LOC120100722 |
| | ↑ | Zbtb16 | ↑ | Map3k6 | ↓↓ | Acta1 | ↓ | Cytl1 | ↑ | Gabrd | ↓ | LOC108348619 |
| | ↑ | Mt1 | ↑ | Fibin | ↑ | Per3 | ↓ | LOC120096903 | ↓↓ | Lbp | ↓ | Igfbp3 |
| | ↑ | Mt2A | ↑ | Plin4 | ↓ | Igfbp3 | ↓ | Pappa2 | ↓↓ | LOC100910099 | ↓↓ | Mpz |
| | ↑ | Pcdh20 | ↑ | Tmem52 | ↓ | LOC100360413 | ↑ | LOC688459 | ↓ | Clec1a | ↓ | LOC108350691 |
| | ↑ | Azin1 | ↑ | Iqgap2 | ↑ | LOC120094629 | ↑ | LOC103694018 | ↓ | Slco2a1 | ↓ | S100a9 |
| | ↑ | Gpatch4 | ↑ | Slc19a3 | ↓ | LOC102552770 | ↑ | Frem1 | ↓ | Ptgdr2 | ↓ | Ttn |
| | ↑ | Cdkn1a | ↑ | Tinagl1 | ↑↑ | Col28a1 | ↑ | RT1-T24-4 | ↓ | Col14a1 | ↓ | Sypl2 |
| **Muscle** | ↑↑ | Nr4a3 | ↑ | Map1b | ↓ | Arntl | ↓ | Npas2 | ↑ | Hyal1 | ↑ | Myh1 |
| | ↑↑ | Zbtb16 | ↑ | Nt5dc3 | ↑ | Per2 | ↑ | Slc41a3 | ↓ | RT1-Ba | ↓ | Tmeff2 |
| | ↑↑ | Ca12 | ↑↑ | Mt1 | ↑ | Per3 | ↓ | LOC120103476 | ↓↓ | Cd8a | ↓ | RT1-Bb |
| | ↑↑ | Fkbp5 | ↑ | Midn | ↓ | Nfil3 | ↓ | Plek | ↓ | Dennd1c | ↓ | Gpsm3 |
| | ↑ | Foxo1 | ↑↑ | Pdk4 | ↑ | Dbp | ↑ | Sntb1 | ↓ | LOC102551352 | ↓ | Irs3 |
| | ↑ | Gpr157 | ↑ | Retreg1 | ↓ | Lrrc30 | ↓ | LOC103693683 | ↓ | Ccr2 | ↓ | Angptl4 |
| | ↑ | Fmo2 | ↑ | Glul | ↑ | Rusc2 | ↑ | Tle4 | ↓ | Ptges | ↑ | Ppm1e |
| | ↑↑ | Adh1 | ↑ | Nr6a1 | ↓ | Auts2l1 | ↑ | Ppp1r3b | ↑ | Mn1 | ↑ | Tst |
| | ↑↑ | Noct | ↑ | Pde4d | ↑ | Usp2 | ↑ | Abr | ↓ | Lck | ↓ | Cyp2j3 |
| | ↑ | Chd7 | ↑↑ | LOC102547513 | ↓ | Srebf1 | ↑ | Ciart | ↓ | Pi16 | ↑ | Bhlhe40 |

*legend:* ↑ increased gene expression relative to SPS-unexposed control rats;
↓ decreased gene expression relative to SPS-unexposed control rats;
↓↓/↑↑ $|\log_2(\text{foldchange})| \geq 2.5$;
↓↓↓/↑↑↑ $|\log_2(\text{foldchange})| \geq 5$

***Supplementary Figure 4.*** The top 20 DEGs at each post-SPS timepoint (based on $|\log_2$*(fold change)*$| \times -\log(p)$ product value) in left hippocampus, right hippocampus, spinal cord, and muscle are listed along with an arrow depicting whether gene expression increased (blue) or decreased (yellow) and the relative magnitude of this decrease (as defined in the legend).

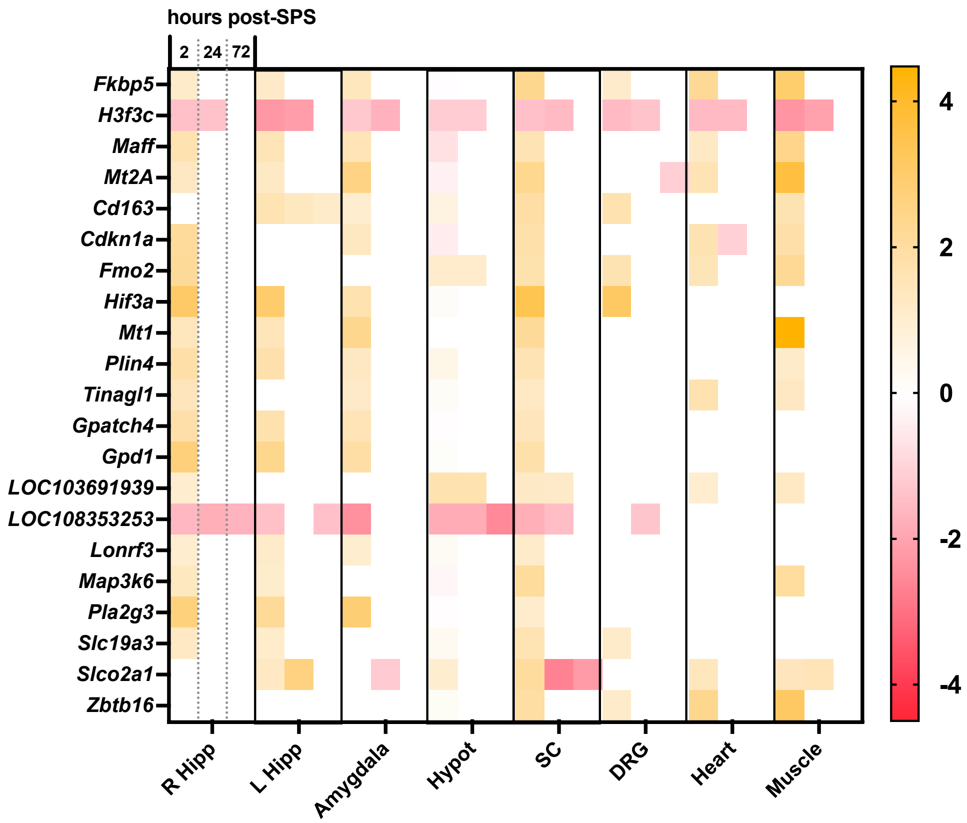


***Supplementary Figure 5.*** Summary of statistically significant differentially expressed genes (DEGs) across multiple timepoints and tissues. The 20 DEGs that were identified as statistically significant ($p \leq 0.05$ and $|\log_2(\textit{fold change})| \geq 1$) across five or more tissues at the 2-hours post-SPS timepoint are shown. Cell color is representative of each gene's $\log_2(\textit{fold change})$ with positive values in yellow and negative values in red. *Abbreviations: R Hipp- Right Hippocampus, L Hipp- Left Hippocampus, Hypot-Hypothalamus, SC-Spinal Cord, DRG-dorsal root ganglion.*

***Supplementary Figure 6.*** Common differentially expressed genes (DEGs) across time and tissues. In *A-K*, counts of statistically significant DEGs are shown in table and Venn diagram cells. In the table view, the count of overlapping DEGs between the tissues and timepoints indicated in the column and row headers is shown and colored according to magnitude. The total number of DEGs is listed in the column and row headers. In the Venn view, the count of overlapping genes between all three timepoints is shown as well as the list of specific genes common to all compared groups. Venn diagrams were not generated for *S6 A-C* due to the number of groups compared.

*S6. A. 2 Hours Post-SPS*
*S6. B 24 Hours Post-SPS*
*S6. C 72 Hours Post-SPS*
*S6. D Right Hippocampus*
*S6. E. Left Hippocampus*
*S6. F. Amygdala*
*S6. G. Hypothalamus*
*S6. H. Spinal Cord*
*S6. I. Dorsal Root Ganglia*
*S6. J. Muscle*
*S6. K. Heart*

***S6. A.*** *2 Hours Post-SPS*

| | Heart (276 DEGs) | DRG (100 DEGs) | Spine (136 DEGs) | Hypothalamus (227 DEGs) | Amygdala (204 DEGs) | Left Hippocampus (146 DEGs) | Right Hippocampus (221 DEGs) |
|---|---|---|---|---|---|---|---|
| Muscle (398 DEGs) | 49 | 15 | 22 | 20 | 14 | 12 | 25 |
| Heart (276 DEGs) | | 12 | 13 | 17 | 13 | 9 | 16 |
| DRG (100 DEGs) | | | 15 | 15 | 10 | 6 | 10 |
| Spine (136 DEGs) | | | | 43 | 20 | 26 | 28 |
| Hypothalamus (227 DEGs) | | | | | 27 | 35 | 40 |
| Amygdala (204 DEGs) | | | | | | 27 | 34 |
| Left Hippocampus (146 DEGs) | | | | | | | 42 |

***S6. B.*** *24 Hours Post-SPS*

| | Heart (82 DEGs) | DRG (97 DEGs) | Spine (78 DEGs) | Hypothalamus (72 DEGs) | Amygdala (178 DEGs) | Left Hippocampus (158 DEGs) | Right Hippocampus (355 DEGs) |
|---|---|---|---|---|---|---|---|
| Muscle (174 DEGs) | 14 | 9 | 4 | 6 | 8 | 4 | 10 |
| Heart (82 DEGs) | | 9 | 3 | 2 | 5 | 2 | 3 |
| DRG (97 DEGs) | | | 8 | 5 | 8 | 7 | 7 |
| Spine (78 DEGs) | | | | 6 | 10 | 8 | 10 |
| Hypothalamus (72 DEGs) | | | | | 5 | 6 | 9 |
| Amygdala (178 DEGs) | | | | | | 16 | 35 |
| Left Hippocampus (158 DEGs) | | | | | | | 24 |

***S6. C.*** *72 Hours Post-SPS*

| | Heart (90 DEGs) | DRG (79 DEGs) | Spine (93 DEGs) | Hypothalam us (44 DEGs) | Amygdala (79 DEGs) | Left Hippoca mpus (254 DEGs) | R Hippocam pus (117 DEGs) |
|---|---|---|---|---|---|---|---|
| Muscle (189 DEGs) | 3 | 8 | 4 | 1 | 6 | 11 | 3 |
| Heart (90 DEGs) | | 4 | 3 | 1 | 1 | 2 | 1 |
| DRG (79 DEGs) | | | 1 | 3 | 3 | 4 | 3 |
| Spine (93 DEGs) | | | | 1 | 2 | 13 | 4 |
| Hypothalam us (44 DEGs) | | | | | 4 | 8 | 2 |
| Amygdala (79 DEGs) | | | | | | 12 | 7 |
| Left Hippoca mpus (254 DEGs) | | | | | | | 18 |

*S6. D.*

### Right Hippocampus

| | R Hippocampus 24 Hours (355 DEGs) | R Hippocampus 72 Hours (117 DEGs) |
|---|---|---|
| R Hippocampus 2 Hours (221 DEGs) | 42 | 13 |
| R Hippocampus 24 Hours (355 DEGs) | | 79 |

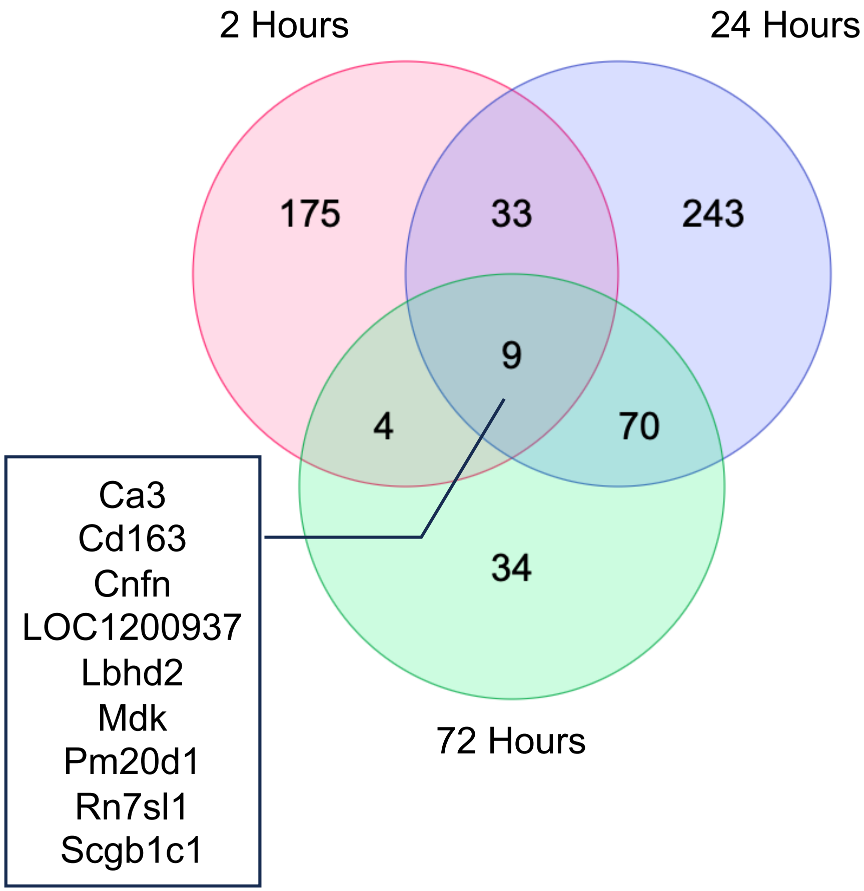

*S6. E.*

### Left Hippocampus

| | L Hippocampus 24 Hours (158 DEGs) | L Hippocampus 72 Hours (254 DEGs) |
|---|---|---|
| L Hippocampus 2 Hours (146 DEGs) | 20 | 8 |
| L Hippocampus 24 Hours (158 DEGs) | | 43 |

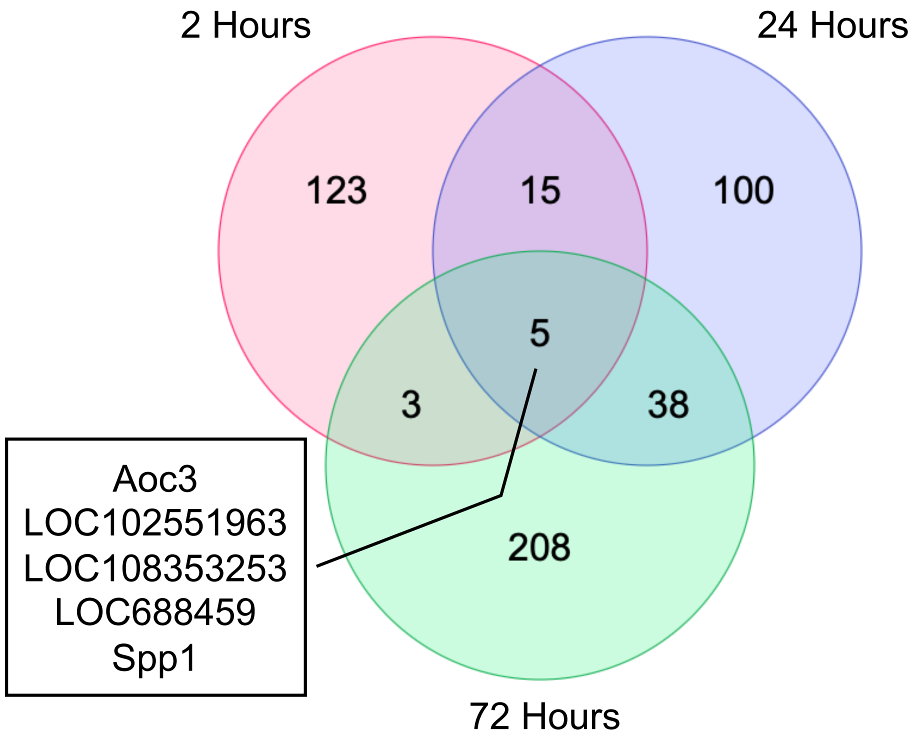

*S6. F.*

### Amygdala

| | Amygdala 24 Hours (178 DEGs) | Amygdala 72 Hours (79 DEGs) |
|---|---|---|
| Amygdala 2 Hours (204 DEGs) | 20 | 14 |
| Amygdala 24 Hours (178 DEGs) | | 31 |

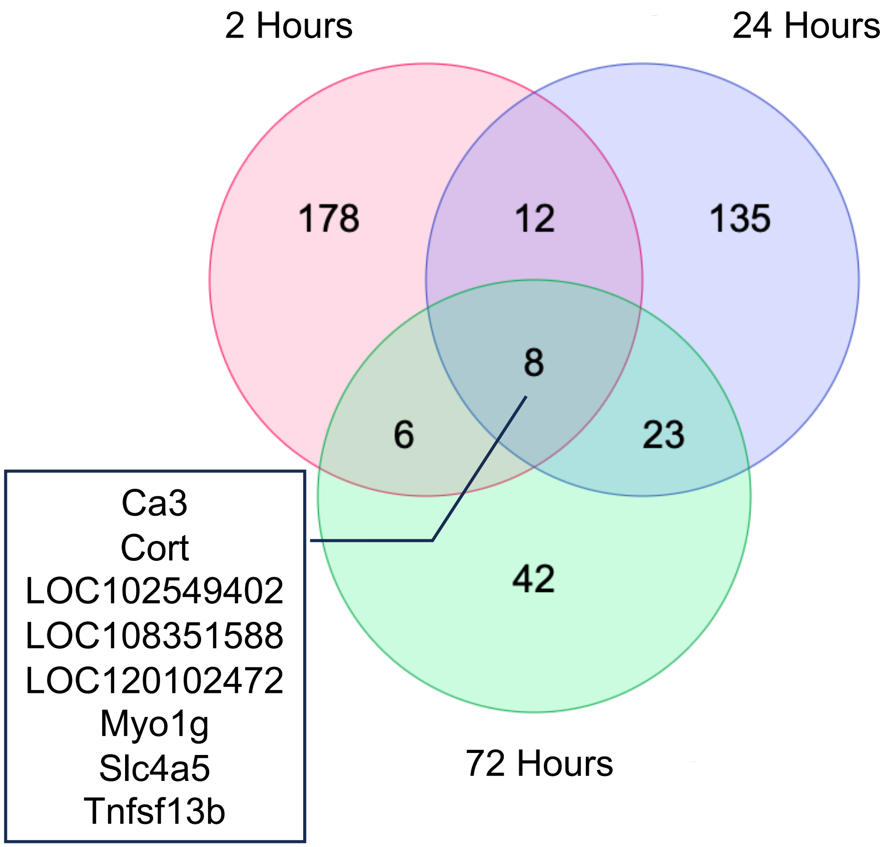

*S6. G.*

### Hypothalamus

| | Hypothalamus 24 Hours (72 DEGs) | Hypothalamus 72 Hours (44 DEGs) |
|---|---|---|
| Hypothalamus 2 Hours (227 DEGs) | 24 | 12 |
| Hypothalamus 24 Hours (72 DEGs) | | 9 |

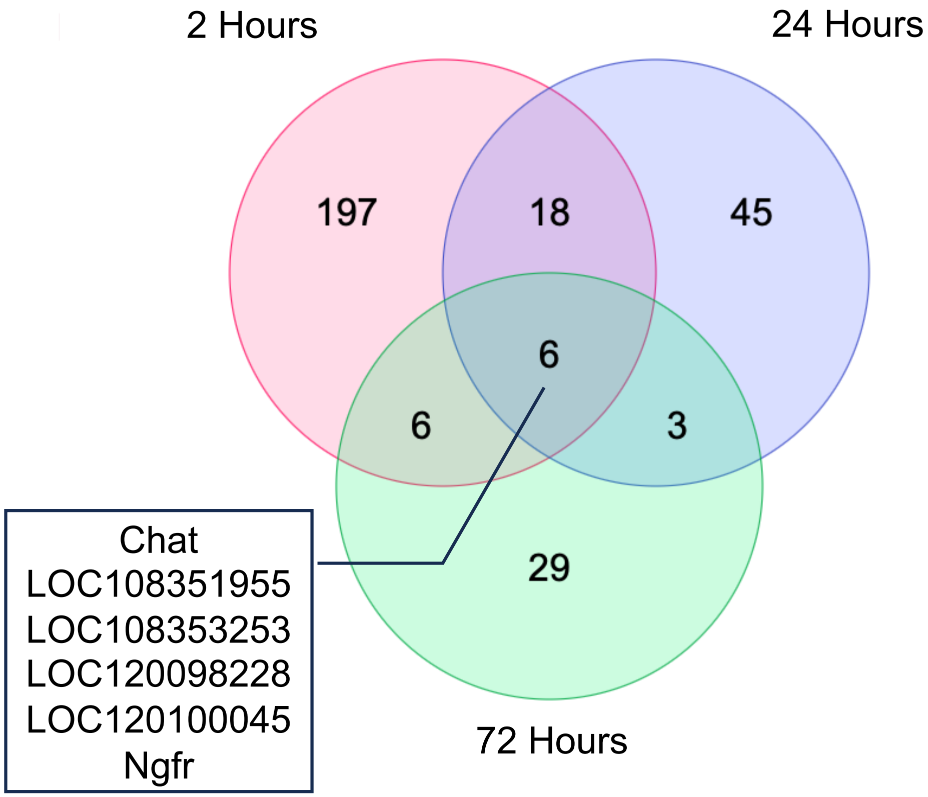

*S6. H.*

#### Spinal Cord

| | Spinal Cord 24 Hours (78 DEGs) | Spinal Cord 72 Hours (93 DEGs) |
|---|---|---|
| Spinal Cord 2 Hours (136 DEGs) | 16 | 15 |
| Spinal Cord 24 Hours (78 DEGs) | | 21 |

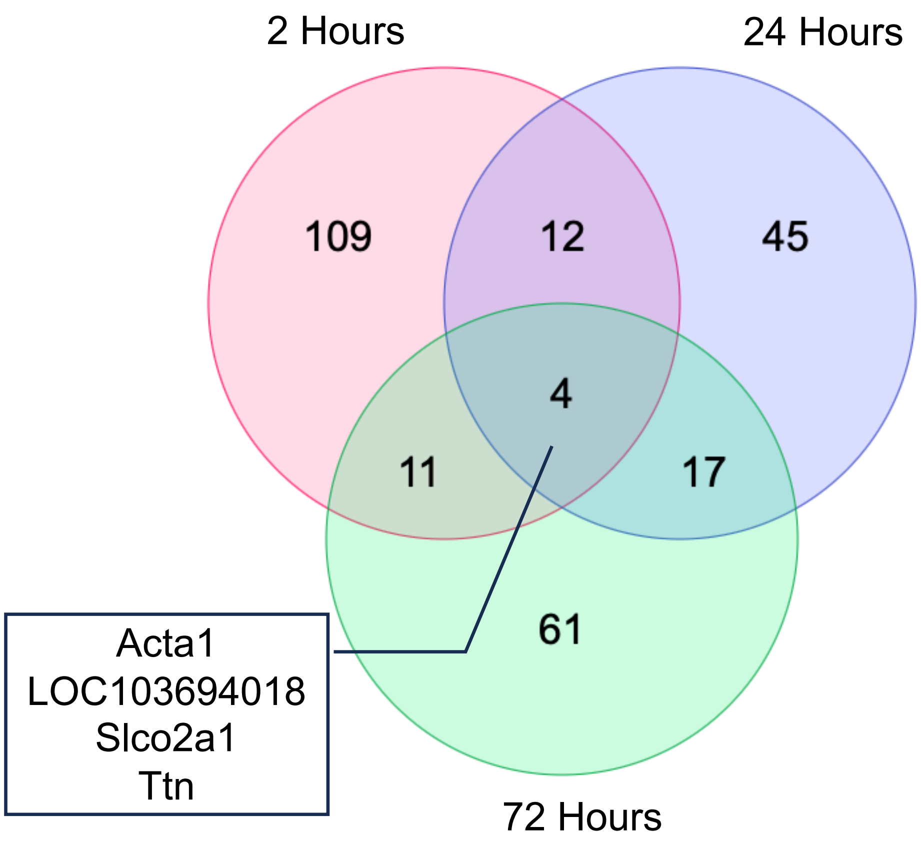

***S6. I.***

### DRG

| | DRG 24 Hours (97 DEGs) | DRG 72 Hours (79 DEGs) |
|---|---|---|
| DRG 2 Hours (100 DEGs) | 15 | 20 |
| DRG 24 Hours (97 DEGs) | | 20 |

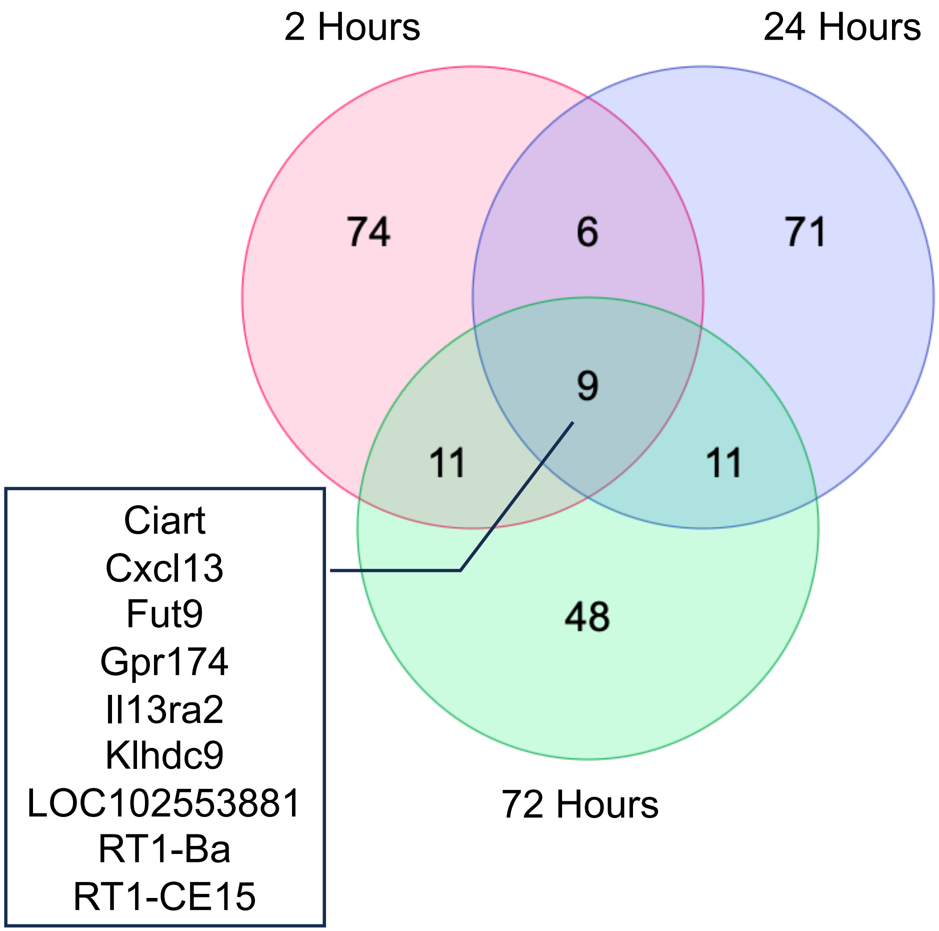

*S6. J.*

### Muscle

| | Muscle 24 Hours (174 DEGs) | Muscle 72 Hours (189 DEGs) |
|---|---|---|
| Muscle 2 Hours (398 DEGs) | 28 | 21 |
| Muscle 24 Hours (174 DEGs) | | 28 |

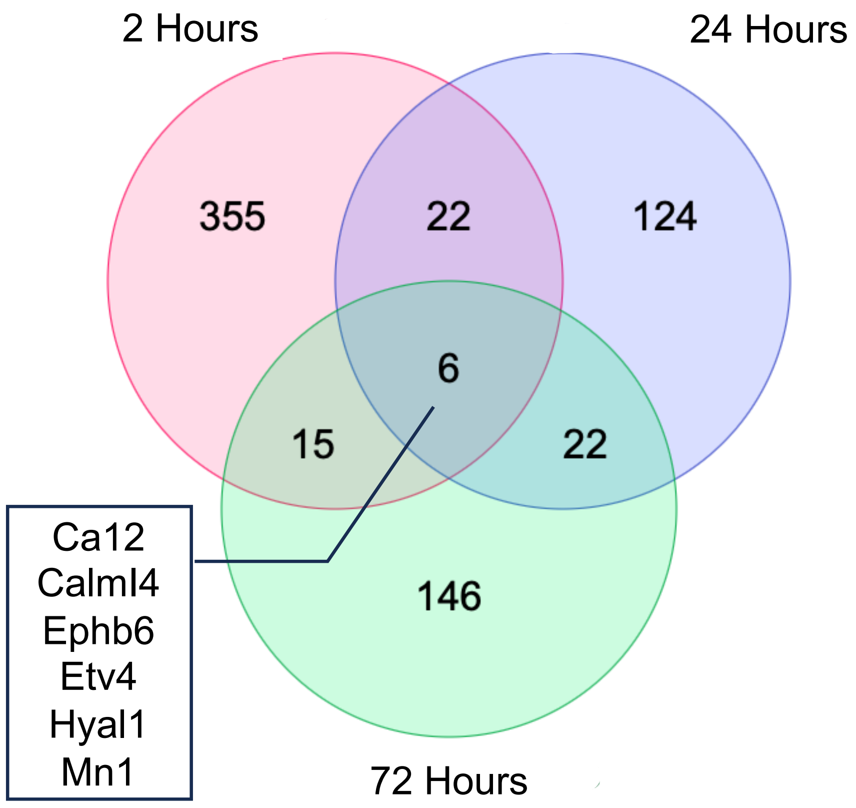

*S6. K.*

### Heart

| | Heart 24 Hours (82 DEGs) | Heart 72 Hours (90 DEGs) |
|---|---|---|
| Heart 2 Hours (276 DEGs) | 20 | 17 |
| Heart 24 Hours (82 DEGs) | | 17 |

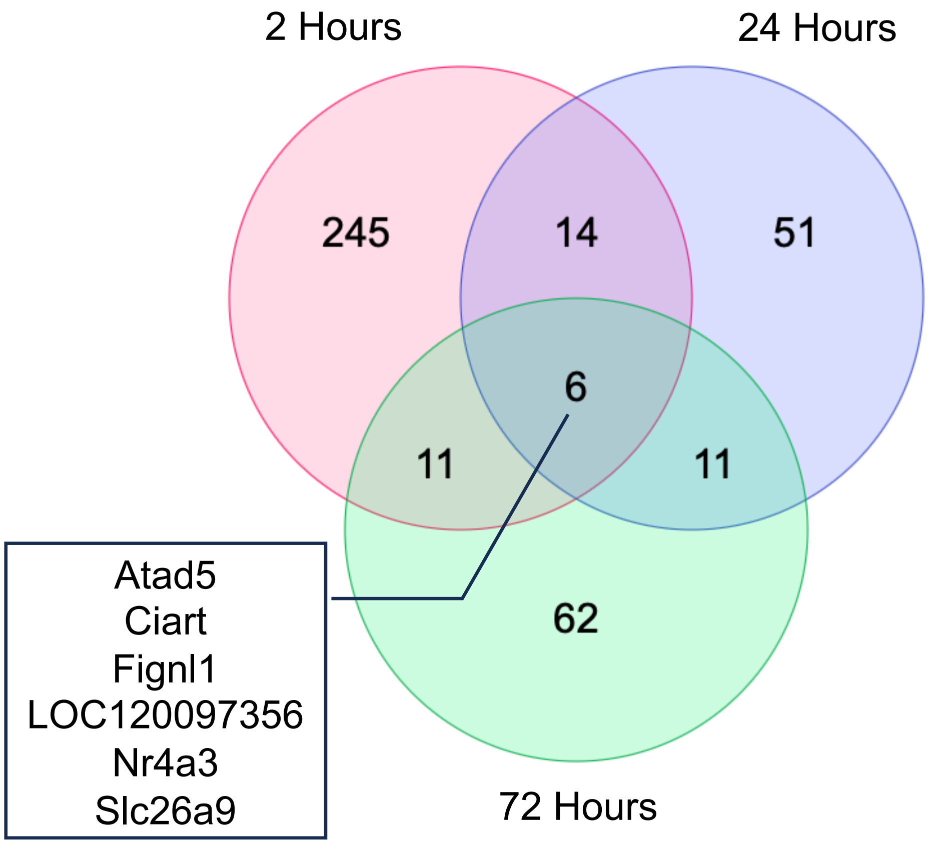

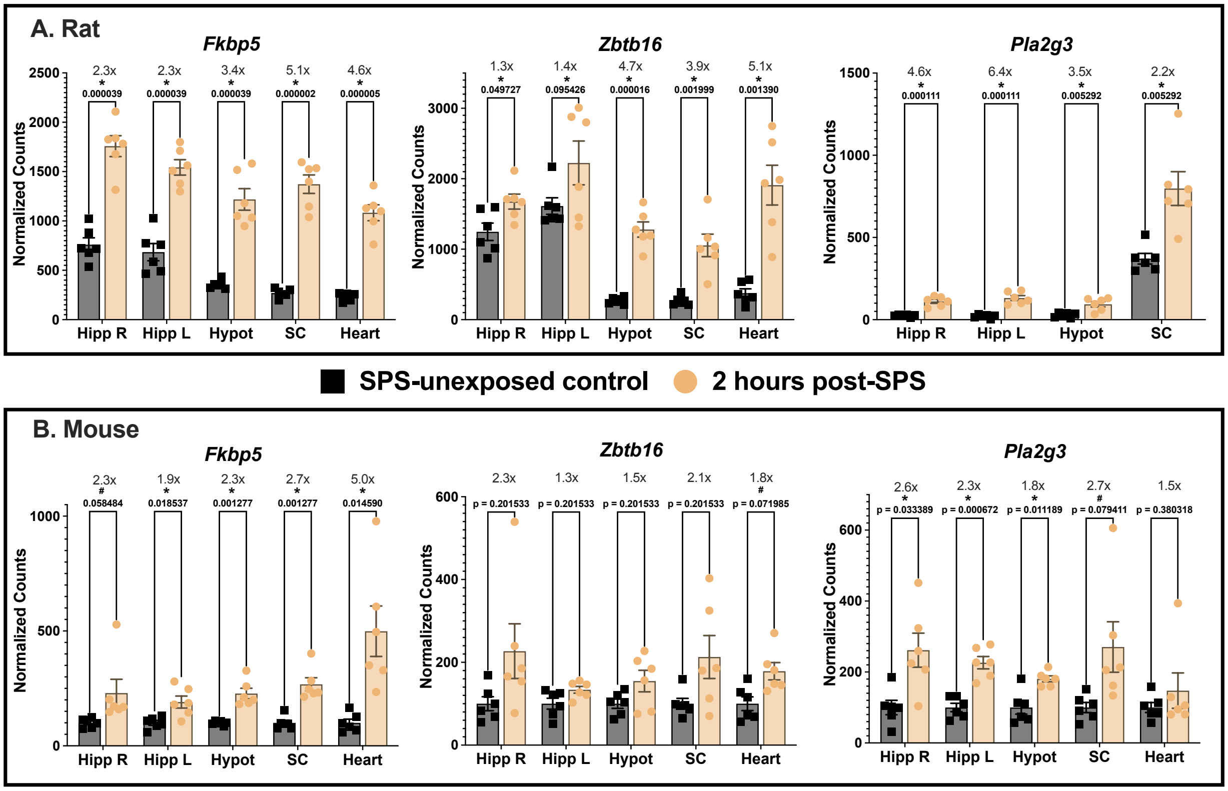


***Supplementary Figure 7.*** Replication of three representative DEGs (*Fkbp5*, *Zbtb16*, and *Pla2g3*) at 2-hours following SPS across rats and mice. **A)** DESeq2 normalized sequencing read counts in rats is shown for transcripts *Fkbp5*, *Zbtb16*, and *Pla2g3* in SPS-unexposed (n=6) and 2-hour post-SPS groups (n=6). After DESeq2 filtering, *Pla2g3* was not detectable in heart tissues, so no data is presented. **B)** Expression patterns of these transcripts following SPS were validated across labs, in mice, and across RNA detection techniques using RT-qPCR. Bars represent Mean ± SEM. **$p<0.05$, $^{\#}p<0.1$. Abbreviations: Hipp-hippocampus, Hypot-hypothalamus, SC-spinal cord, DEG-differentially expressed gene, SPS-single prolonged stress, mSPS-mouse SPS model, qPCR-quantitative polymerase chain reaction.*

| | KEGG Pathway | Product | FDR | Enrichment Score | KEGG Pathway | Product | FDR | Enrichment Score | KEGG Pathway | Product | FDR | Enrichment Score |
|---|---|---|---|---|---|---|---|---|---|---|---|---|
| Right Hippocar | Protein digestion and absorption | 13.84 | 0.00 | 4.59 | Protein digestion and absorption | 12.19 | 0.00 | 4.68 | Protein digestion and absorption | 20.84 | 0.00 | 7.74 |
| | Thyroid hormone signaling | 12.56 | 0.00 | 4.17 | ECM-receptor interaction | 7.18 | 0.02 | 4.22 | AGE-RAGE signaling in diabetic | 16.18 | 0.00 | 6.98 |
| | Proteoglycans in cancer | 7.62 | 0.00 | 3.06 | GABAergic synapse | 7.09 | 0.02 | 4.17 | Platelet activation | 15.41 | 0.00 | 6.42 |
| | Human papillomavirus infection | 7.42 | 0.00 | 2.64 | Human papillomavirus infection | 4.95 | 0.01 | 2.56 | Proteoglycans in cancer | 8.64 | 0.01 | 4.44 |
| | TGF-beta signaling pathway | 7.20 | 0.01 | 3.83 | Platelet activation | 4.04 | 0.06 | 3.24 | ECM-receptor interaction | 7.41 | 0.05 | 5.81 |
| Left Hippocam | Nitrogen metabolism | 37.72 | 0.01 | 17.04 | Circadian entrainment | 40.98 | 0.00 | 9.13 | Protein digestion and absorption | 59.72 | 0.00 | 8.39 |
| | Thyroid hormone signaling | 11.92 | 0.01 | 5.39 | Glycerolipid metabolism | 18.16 | 0.01 | 8.04 | ECM-receptor interaction | 24.87 | 0.00 | 6.71 |
| | Nicotinate and nicotinamide | 11.21 | 0.04 | 8.04 | Cholinergic synapse | 16.35 | 0.00 | 6.39 | PI3K-Akt signaling pathway | 18.72 | 0.00 | 3.81 |
| | Ras signaling pathway | 8.51 | 0.01 | 3.84 | Apelin signaling pathway | 14.95 | 0.00 | 5.84 | Hypertrophic cardiomyopathy | 10.34 | 0.01 | 5.07 |
| | Calcium signaling pathway | 8.29 | 0.01 | 3.74 | GABAergic synapse | 10.64 | 0.01 | 5.80 | Human papillomavirus infection | 9.04 | 0.00 | 3.14 |
| Amygdala | alpha-Linolenic acid metabolism | 8.76 | 0.06 | 7.32 | ECM-receptor interaction | 61.05 | 0.00 | 7.47 | Pertussis | 19.75 | 0.00 | 7.46 |
| | Linoleic acid metabolism | 6.92 | 0.06 | 5.78 | Focal adhesion | 21.23 | 0.00 | 3.96 | Sulfur metabolism | 18.77 | 0.06 | 15.35 |
| | Non-alcoholic fatty liver disease | 5.44 | 0.03 | 3.47 | PI3K-Akt signaling pathway | 17.67 | 0.00 | 3.22 | Staphylococcus aureus infection | 18.73 | 0.00 | 6.75 |
| | Ether lipid metabolism | 4.90 | 0.09 | 4.77 | Proteoglycans in cancer | 17.36 | 0.00 | 3.70 | Lipid and atherosclerosis | 16.74 | 0.00 | 5.09 |
| | Cocaine addiction | 4.79 | 0.09 | 4.67 | Protein digestion and absorption | 15.44 | 0.00 | 4.49 | Phagosome | 16.10 | 0.00 | 5.24 |
| Hypothalamus | ECM-receptor interaction | 18.69 | 0.00 | 4.40 | Circadian rhythm | 114.17 | 0.00 | 24.85 | None | | | |
| | FoxO signaling pathway | 12.58 | 0.00 | 3.39 | Viral myocarditis | 9.07 | 0.07 | 8.05 | | | | |
| | Insulin resistance | 11.61 | 0.00 | 3.46 | Phagosome | 6.19 | 0.07 | 5.25 | | | | |
| | Human papillomavirus infection | 11.44 | 0.00 | 2.57 | Epstein-Barr virus infection | 5.73 | 0.06 | 4.81 | | | | |
| | Proteoglycans in cancer | 11.40 | 0.00 | 2.92 | | | | | | | | |
| Spinal Cord | Proteoglycans in cancer | 9.36 | 0.01 | 4.09 | Circadian rhythm | 16.20 | 0.10 | 16.05 | Protein digestion and absorption | 63.98 | 0.00 | 11.61 |
| | Small cell lung cancer | 7.06 | 0.04 | 4.89 | | | | | AGE-RAGE signaling in diabetic | 22.48 | 0.00 | 8.37 |
| | PI3K-Akt signaling pathway | 6.07 | 0.01 | 3.06 | | | | | ECM-receptor interaction | 18.90 | 0.01 | 8.36 |
| | Thyroid hormone signaling | 6.00 | 0.04 | 4.16 | | | | | p53 signaling pathway | 16.90 | 0.01 | 8.34 |
| | ECM-receptor interaction | 5.42 | 0.06 | 4.44 | | | | | Breast cancer | 11.59 | 0.01 | 5.72 |
| DRG | Transcriptional misregulation in | 16.70 | 0.00 | 5.63 | Circadian rhythm | 58.19 | 0.00 | 19.30 | Asthma | 57.12 | 0.01 | 29.94 |
| | Asthma | 12.41 | 0.07 | 10.92 | Cytokine-cytokine receptor interaction | 13.15 | 0.00 | 5.09 | Circadian rhythm | 41.82 | 0.01 | 22.90 |
| | Endocrine resistance | 10.13 | 0.02 | 6.17 | Viral protein interaction with cytokine and | 13.15 | 0.02 | 8.00 | Vitamin digestion and absorption | 29.38 | 0.04 | 20.76 |
| | Breast cancer | 9.99 | 0.01 | 5.22 | Staphylococcus aureus infection | 11.85 | 0.02 | 7.21 | Viral myocarditis | 28.29 | 0.01 | 14.83 |
| | Melanoma | 9.63 | 0.04 | 6.67 | Viral myocarditis | 9.80 | 0.05 | 7.50 | Graft-versus-host disease | 27.78 | 0.02 | 16.56 |
| Muscle | Focal adhesion | 14.29 | 0.00 | 2.93 | Circadian rhythm | 54.82 | 0.00 | 10.50 | Asthma | 52.78 | 0.00 | 12.31 |
| | Small cell lung cancer | 13.25 | 0.00 | 3.59 | Insulin resistance | 8.39 | 0.01 | 3.97 | Systemic lupus erythematosus | 42.63 | 0.00 | 7.36 |
| | FoxO signaling pathway | 10.82 | 0.00 | 3.01 | Starch and sucrose metabolism | 6.67 | 0.08 | 5.95 | Staphylococcus aureus infection | 40.76 | 0.00 | 7.04 |
| | EGFR tyrosine kinase inhibitor | 10.34 | 0.00 | 3.45 | Cysteine and methionine metabolism | 4.62 | 0.08 | 4.28 | Pertussis | 40.53 | 0.00 | 7.62 |
| | TGF-beta signaling pathway | 10.05 | 0.00 | 3.25 | Insulin signaling pathway | 3.28 | 0.08 | 2.93 | Antigen processing and | 32.22 | 0.00 | 7.09 |
| Heart | Circadian rhythm | 18.35 | 0.00 | 6.08 | Circadian rhythm | 228.23 | 0.00 | 25.25 | Circadian rhythm | 136.47 | 0.00 | 21.65 |
| | Apoptosis | 7.65 | 0.00 | 3.00 | Malaria | 14.91 | 0.02 | 8.76 | | | | |
| | Glycerophospholipid | 7.37 | 0.00 | 3.20 | cGMP-PKG signaling pathway | 13.43 | 0.00 | 5.24 | | | | |
| | Small cell lung cancer | 7.04 | 0.01 | 3.22 | Regulation of lipolysis in adipocytes | 12.41 | 0.03 | 7.81 | | | | |
| | Toxoplasmosis | 5.94 | 0.01 | 2.87 | Circadian entrainment | 8.49 | 0.03 | 5.66 | | | | |

**Legend**

| | | | | |
|---|---|---|---|---|
| Stress Signaling | Circulatory System | Neuro-Specific Signaling Processes | Metabolism | General Cellular Processes |
| Cancer | Immune/Inflammatory Signaling | Genetic/Epigenetic Machinery | Circadian | Other |

***Supplementary Figure 8.*** Top 5 KEGG Ontologies by post-SPS Timepoint and Tissue. Kyoto Encyclopedia of Genes and Genomes (KEGG) pathways enriched in DEGs for each timepoint and tissue (FDR $q \leq 0.1$) were sorted in descending order based on their product score (i.e. $|\log_2(\textit{fold change})| \times -\log(p)$). Each KEGG pathway is colored according to their category designation and shown alongside their product value, FDR $q$ value, and enrichment score.
